# Operator-valued free probability applied to recurrent neural networks with heterogeneity in local and global circuit structures

**DOI:** 10.64898/2026.10.08.757745

**Authors:** Juliet Jiang, Gengshuo John Tian, Brent Doiron

## Abstract

Random connectivity is a common assumption in models of recurrent neural networks, yet biological circuits combine heterogeneous synaptic connections with organization into distinct anatomical regions and functional groups. How does this global structure interact with cellular and local circuit heterogeneity to shape the dimensionality of population activity? In our work, we use operator-valued free probability to derive the covariance spectrum and effective dimension of recurrent networks with heterogeneous gains and block-structured random connectivity. This framework handles the noncommuting matrix operations through which cellular and circuit heterogeneity jointly determine population variability. We show that the linear relationship between effective dimension and firing rate uniformity persists despite global connectivity structure. At fixed total connectivity variance and with identical firing rate distributions across areas, concentrating variance within one area decreases dimension, whereas concentrating it in directed connections between areas increases dimension. Together, these results establish a mechanistic connection between circuit organization and population dimensionality, and demonstrate how operator-valued free probability unifies cellular, local, and global sources of heterogeneity in recurrent networks.

## I. INTRODUCTION

Biological systems are generally complex, with a plethora of components and factors to consider [1]. External inputs to a system and the internal interactions of its components combine to determine output behavior. In some cases complex activity is simply *inherited* from diverse external inputs, and the system can be well described by a passive filter (Figure 1a). A classical example in the visual system is the Hermann grid illusion, which can be understood by modeling neurons as linear filters of the stimulus [2, 3]. Other examples include the cochlea which acts as a wavelet transform of the acoustic signal [4]; the spatial distribution of plankton due to heterogeneous inertial properties in response to oceanic flows [5]; and the functional landscape of microbial communities, which can often be well-predicted simply by the presence or absence of species without modeling their interactions [6]. Alternatively, complex dynamics can *emerge* from interactions within the system (Figure 1b). For example, visual hallucinations arise from recurrent interactions between neurons without any visual stimuli [7, 8]. Similarly, the dynamical patterns in bird flocking [9], morphogenesis [10], and central pattern generators [11] all self-organize from internal dynamics within the system.

**Figure 1.**
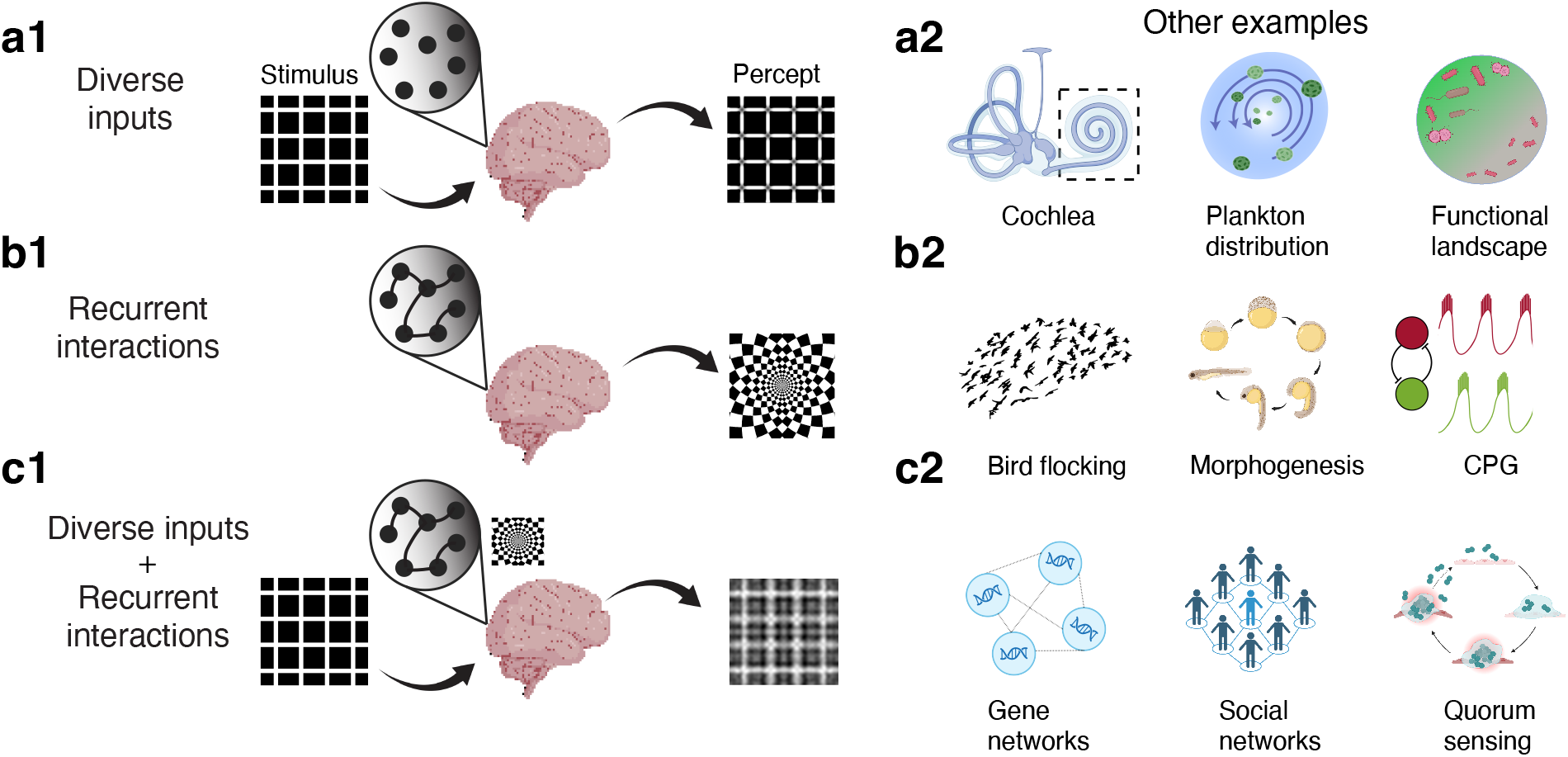
**(a-c)** Three types of models for how biological systems produce complex outputs. **a)** The system acts as a passive filter of the external input. **a1)** The Hermann grid visual illusion, which can be explained by neurons responding linearly to the grid image [2, 3]. **a2)** Three other examples: the cochlea as a wavelet transform machine [4], plankton distribution due to different sizes responding differently to the flow [5], and the functional landscape of microbial communities as a linear or quadratic function of the presence of species [6]. **b)** Patterns spontaneously emerge from the interactions within the system. **b1)** Visual hallucinations result from recurrent interactions of cortical neurons in a network [7, 8]. **b2)** Three other examples: bird flocking [9], morphogenesis [10], and central pattern generators [11]. **c)** The external input combines with the internal interactions of the system to produce complex behaviors. **c1)** The active processing of the stream of input by the visual system. **c2)** Three other examples: gene networks [12], social networks [13], and quorum sensing bacterial colonies [14].

While these two organization principles for how complex solutions of system activity arise have enjoyed great success, most phenomena result from a combination of diverse external inputs and complex recurrent interactions and should be studied as such (Figure 1c). In the end, it is the active processing of external input that engenders visual cognition [15]. Incorporating this complexity is especially crucial when studying systems like gene networks [12], social networks [13], and quorum sensing bacterial colonies [14] where the system functions in response to changes in the environment through the interactions of its components. The central goal of this study is to put forth a coherent theory for how the heterogeneities of inputs and interactions combine to determine system output. We focus our efforts on the nervous system, yet our approach can be applied in a wide array of disciplines.

While our goal is simply stated, it becomes especially difficult when the system consists of a large number of highly diverse and interacting components, making the problem irreducibly high-dimensional. This problem is particularly critical in the study of neuronal circuits, in which thousands or even millions of neurons integrate long-range synaptic inputs (feedforward and feedback) through rich recurrent dynamics to produce computationally meaningful responses. To date, while simulations and fitted models have incorporated high degrees of detail [16, 17], theoretical understanding has mostly been based on models that consider only one aspect of heterogeneity in the network [18–22].

Recently, we advanced this research program by studying how random, yet unstructured, recurrent connectivity combines with diverse external inputs to produce complex network responses [23]. We used (operator-valued) free probability theory [24] to derive that the effective dimension of the response covariance matrix (spread of its eigenvalues) depends on the product of the uniformity of the firing rates (proxy for the external drive) and the variability of recurrent weights (internal interactions). The prediction that the effective dimension is linearly related to the uniformity of firing rates was tested in various neuronal datasets across the cortices of mice, monkeys and humans. Further, we showed that when animals are in a state of heightened processing (i.e., performing a task), the uniformity of firing rates decreases and subsequently the effective dimension decreases, both acting to improve stimulus discrimination.

While this agreement between experiment and theory is satisfying, it also raises deeper issues. While unstructured random wiring is a convenient mathematical construct to explore network activity [19, 21, 25–28], it ignores a wealth of experimental work that catalogs hierarchical wiring across brain regions [29, 30], as well as functional and circuit assembly structures within a brain region [31–33]. An open question is then: does the linear relationship between the effective dimension of population activity and the uniformity of firing rates extend to networks with connectivity structures that are much better mimics of real neuronal networks? In this study, we explore this central question and expand our free probability treatment of recurrent networks to include a block structure in connectivity to model distinct anatomical or functional sub-groups of neurons.

The paper is organized as follows. Section II presents the basic measures of firing rate uniformity and effective dimension of population covariability that are central to our study. Section III introduces the core circuit model that anchors our work. Sections IV–VI are didactic and present concepts in random matrix theory, non-commutative free probability, and its extension to non-commutative random variables equipped with block structure. These techniques form the mathematical back-bone of the later sections, and we devote effort in their presentation because they remain non-standard for the physics community. In Section VII we follow our past study [23] and use free probability techniques to relate the uniformity of firing rates to the effective dimension of covariability. Section VIII then asks how this relationship changes under a global structure that groups neurons into two ‘areas’. Sections VIII A-VIII C break these differences into three main types: when the recurrent connections within an area, the projections from one area to another, or all connections and projections from/to an area are differentially diverse. We also consider other cases where these differences are symmetric vs. asymmetric across areas. Finally, Section IX questions how the geometry of neuronal responses within an area may be characterized by features of the larger network.

## II. MEASURES OF NEURONAL RESPONSE

It is now routine for experimental neuroscientists to simultaneously record the activity of large numbers of neurons from the brains of behaving animals [34]. Neuropixels probes and microelectrode arrays record spike trains from up to a thousand neurons simultaneously [35– 37], and two-photon calcium imaging methods capture the activity from hundreds to thousands of individual neurons [35, 38], with some recent methods recording hundreds of thousands [39]. In most cases, a dataset contains multiple trials of an experiment, which could involve the presentation of a sensory input, a cognitive task, or a prescribed movement. On trial *t* a spike rate vector 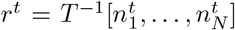 is recorded; 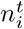 is the number of action potentials (spikes) that neuron *i* ∈ [1, …, *N*] produced on trial *t*, and *T >* 0 is the measurement time window. From *r*^*t*^ we seek to characterize the geometric structure of population responses, and to this end we consider two statistics of population responses. The first is the trial-averaged firing rate *R* = diag[*r*_1_, …, *r*_*N*_], where 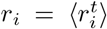 is an expectation across trials. The second is the trial-to-trial covariability matrix *C* with element 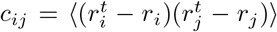. We note that both *R* and *C* are square (*N* ×*N*) Hermitian matrices.

A measure of the dimensionality of population-level response should demonstrate the level of coordination or the degrees of freedom in the population’s collective activity [19, 21, 23, 26, 27, 40, 41]. Indeed, the variability of these high-dimensional neuronal responses can often be sufficiently captured by a subspace of much lower dimension [42–47]. A popular measure for dimensionality is the normalized participation ratio of the covariance matrix *C* [48], which we will label as the *effective dimension*. This measure is governed by the eigenvalues {*λ*_*i*_} of *C*:

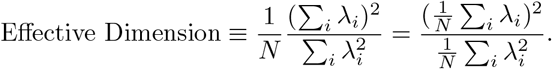

We note that the moments of the eigenvalue distribution of matrix *C* are defined by the expected trace of its powers:

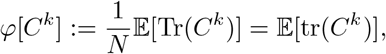

where tr(·) is the normalized trace. Since 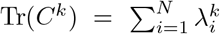 for any positive integer *k*, taking the self-averaging large *N* limit renders the effective dimension a ratio of moments:

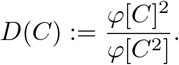

The eigenvalues of *C* depict the amount of population variance in each eigenvector direction given by the activity of neurons. If all the variance is concentrated in one dimension, then *D*(*C*) = 0 in the large *N* limit, while if the variance is evenly spread across all dimensions, then *D*(*C*) = 1 and the response fluctuations are high-dimensional (Figure 2a, left to right).

**Figure 2.**
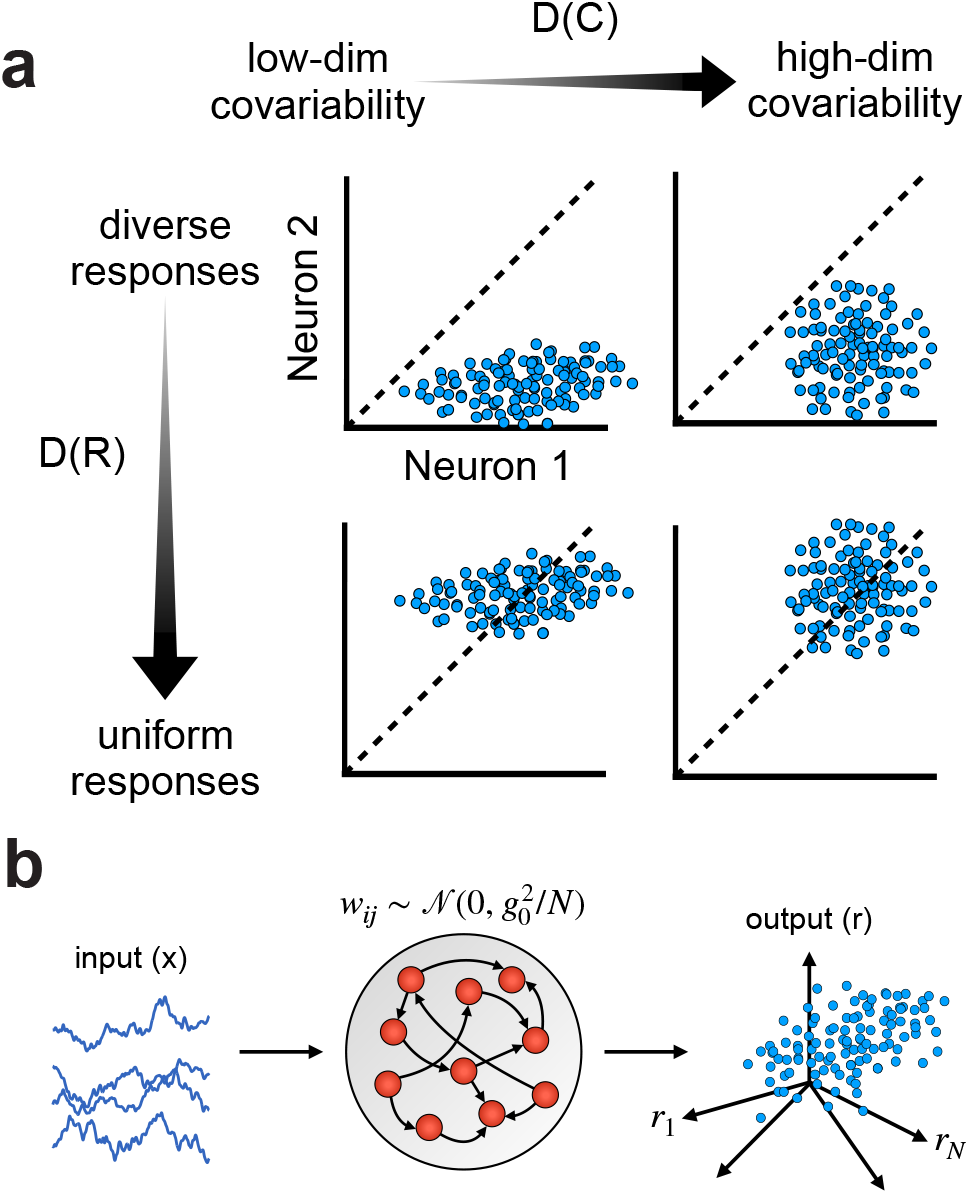
**a)** Projecting the representation onto two dimensions, or two neurons, we show how the properties of example response clouds produce different effective dimension *D*(*C*) and response uniformity *D*(*R*) measures. When the data cloud is elliptical, meaning fluctuations are not evenly distributed across principal directions, *D*(*C*) is low, while for symmetrically sized fluctuations *D*(*C*) is high. When the data cloud is centered off of the unit diagonal, meaning one neuron has a mean firing rate that is higher than the other then *D*(*R*) is low, while if the cloud is centered on the diagonal the firing rates of the neuron pair are similar, and *D*(*R*) is high. The top row has a distribution of neuronal response that has both lower uniformity in firing rates (smaller *D*(*R*)) and lower dimension of covariability (smaller *D*(*C*)). The bottom row shows an example with higher uniformity and higher dimensionality. *R* is sampled from a log-normal distribution (Appendix A). **b)** A neuronal circuit processes external input *x* with its recurrent dynamics through matrix *W*, to produce a neuronal response *r*. The response is represented in *N*-dimensional space, where *N* is the number of neurons in the circuit, and each coordinate is the firing rate of each neuron. Each dot is the response of all *N* neurons for a given realization of input.

We will use the same metric to characterize the *uniformity* of neuronal firing rates:

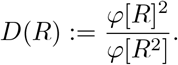

*D*(*R*) is often referred to as the Treves-Rolls measure [49–51], and is a measure of the spread (or diversity) of firing rates. Larger values of *D*(*R*) indicate greater uniformity in firing rates, whereas smaller values represent greater diversity (Figure 2a, top to bottom).

*D*(*C*) determines the geometry of the response variability and *D*(*R*) describes the placement of the mean. Since *D*(*R*) and *D*(*C*) deal with distinct moments of a response they, in principle, have no explicit relation to one another (Figure 2a). However, as we will explore in later sections, circuit mechanisms will establish a tight relation between *D*(*C*) and *D*(*R*) [23].

## III. RECURRENT CIRCUIT MODEL OF A NEURONAL NETWORK

Computations in the brain are often mechanistically modeled using networks of neurons that are recurrently connected in a random manner [19, 25, 52–54]. Networks that treat synaptic couplings as random ensembles have been studied extensively: their eigenvalues are used to study spontaneous and evoked activity [55], they model high-dimensional background activity upon which low-dimensional structure can be imposed [56], they serve as initializations for training deep neural networks [57, 58], and more. The outputs of these networks are correlated based on the input-output relations of individual neurons and their synaptic wiring (Figure 2b). We follow past work [19, 23, 59] and derive how recurrent coupling and input-output neuronal transfer combine to determine *C*.

We begin by formalizing both the input, recurrent circuit structure, and outputs of a network of model neurons (Figure 2b). In what follows, given a collection of scalar random variables on realization *t*, 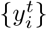, we take a notational convention that we drop the subscript indicator for the vector 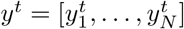, and drop the superscript indicator for expectations 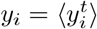.

The *i*^th^ neuron in a recurrent network of *N* neurons with random connectivity has the governing dynamical equation on realization *t* of the random process {*ζ*^*t*^}:

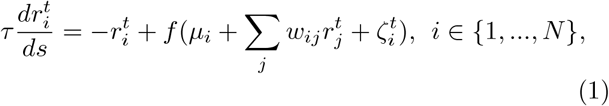

where *s* denotes time, 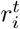 is the response of the neuron, *τ* is the time constant, *µ*_*i*_ is the average external input, *w*_*ij*_ is the synaptic weight from neuron *j* to *i*, and 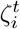 is the trial-to-trial input noise into neuron *i*. Finally, *f* is the neuronal transfer function that maps inputs to output firing rates.

The vectorized form of this equation is:

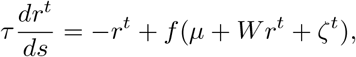

where *W* is the matrix of synaptic weights. Taking the system to steady state, we have *r*_0_ = *f* (*µ* + *Wr*_0_). Assuming noise *ζ* is small, we can linearize around the deterministic steady state *r*_0_ with respect to *ζ*, obtaining

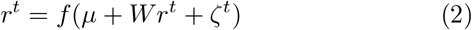

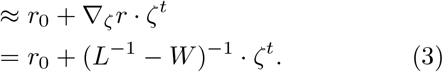

The expression ∇_*ζ*_*r* = (*L*^−1^ − *W*)^−1^ is derived from differentiating Equation (2). Here, *L* is a diagonal matrix of neuronal gains around operating point *r*_0_: *L*_*ii*_ = *f*′(*µ*_*i*_ + ∑_*j*_ *w*_*ij*_*r*_0*j*_).

For the nonlinear transfer function *f*, we assume

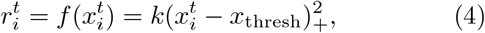

where 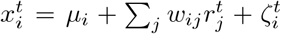, *k* is a constant scaling factor, and *x*_thresh_ is the input threshold [60]. This choice of *f* is such that *L*^2^ ∝ *R*, where *R* = diag[*r*_01_, …, *r*_0*N*_]. By this approximation, the covariance matrix *C* is

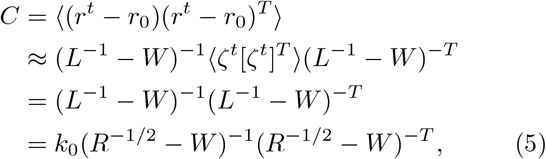

where in the last two steps we assumed the covariance of the input noise ⊗ *ζ*^*t*^[*ζ*^*t*^]^*T*^ ⊗= *I* and pulled all scaling constants (e.g. *k*) into one variable *k*_0_ (with some scaling absorbed into *W*).

In Equation (5) we have derived the covariance matrix *C* in terms of both the weight matrix *W* and firing rate matrix *R*. Past works [19, 21, 26] have taken *R* = *I* and element 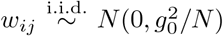, where *g*_0_ sets the effective variability in the strength of recurrent connectivity. Consequently, *C* is a function of just the random matrix *W*, and there are established methods to obtain the eigen-spectrum of *C* (of which the first two moments define *D*(*C*)). In particular, Recanatesi et al. [26] and Hu and Sompolinsky [19] used field theoretic techniques to derive the compact relation:

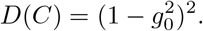

We see that as the variability of connections increases, the effective dimension of activity lowers; this is due to the quenched randomness in *W* promoting certain activity directions over others.

In this study we will consider the case where both *W* and *R* are random matrices. To move forward we will need greater machinery to determine *C*’s eigenspectrum (and hence in turn *D*(*C*)), in part due to the non-commutative nature of *W* and *R*. In the next sections we give a didactic introduction to important topics in random matrix theory and how techniques from free probability theory can be leveraged to compute expectations of *C* when both *W* and *R* are random.

## IV. RANDOM MATRIX THEORY

### A. Eigenvalue distribution

To introduce many of the fundamental concepts in random matrix theory, let us visit the most classic, foundational example: Wigner’s semicircle law [24, 61–64]. Take an *N* × *N* symmetric matrix *A* with independent real Gaussian entries on and above the diagonal. Let the entries have mean 0, variance 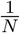 on the off-diagonal, and 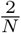 on the diagonal. This matrix is one realization of the Gaussian Orthogonal Ensemble (GOE). Its *N* eigen-values *λ*_1_, …, *λ*_*N*_, counting multiplicity, are encoded in a probability measure on ℝ:

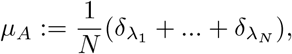

termed the eigenvalue distribution of *A*. The expected empirical distribution E[*µ*_*A*_] is the marginal of the joint distribution of eigenvalues:

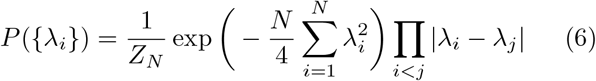

where *Z*_*N*_ is a normalization constant. The two terms in Equation (6) represent distinct aspects of the random matrix *A*. The Gaussian term is inherited from the Gaussian distribution of the elements of *A* and the fact that Tr(*A*^2^) = Tr(Λ^2^), with Λ = diag(*λ*_1_, …, *λ*_*N*_). The term Π_*i<j*_ |*λ*_*i*_ − *λ*_*j*_| ≡ |det(*V*)| is the magnitude of the Vandermonde determinant, where the *i*^th^ row of is [1, *λ*_*i*_, 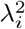, …]. Here, det(*V*) measures the volume of the Jacobian transforming the joint distribution from entries of *A* to eigenvalues in Λ through the diagonalization *A* = *S*Λ*S*^*T*^, where *S* contains the eigenvectors of *A*. Since orthogonal conjugation preserves volume, we can compute the Jacobian with the eigenbasis *I* rather than *S*, while retaining infinitesimal variations of *S* [63]. In effect, this term expresses the fact that a redundant pair of eigenvalues will never exist (*λ*_*i*_ *≠ λ*_*j*_ when *i ≠ j* ∀*i, j*, otherwise *P* ({*λ*_*i*_}) = 0), so eigenvalues instead spread out and never cross.

It is useful to write Equation (6) as:

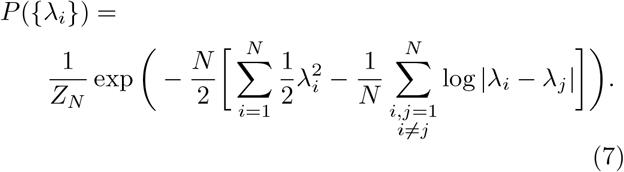

This permits an analogy between the joint distribution of eigenvalues {*λ*_*i*_} and a one-dimensional Coulomb gas of *N* particles, where each eigenvalue represents the location of a particle. The exponent, a finite-*N* energy functional, contains the same two terms as in Equation (6). The first term (Gaussian) analogizes a single particle potential energy from an external field. The second term (Vandermonde determinant) is a pairwise logarithmic potential describing the repulsive forces between charged particles or eigenvalues. The scaled continuous energy functional of measure *µ*_*A*_, or the continuous analogue of the exponent in Equation (7), is then:

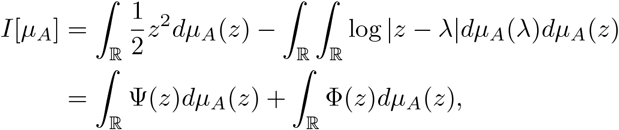

where 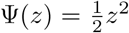 is the external field and Φ is the logarithmic potential

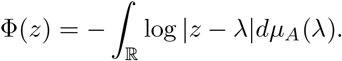

The second term in *I*[*µ*_*A*_], ∫ Φ*dµ*_*A*_, is also called the logarithmic energy from potential theory [65, 66], or the negative of Voiculescu’s free non-commutative entropy from free probability (Figure 3b) [24, 67].

**Figure 3.**
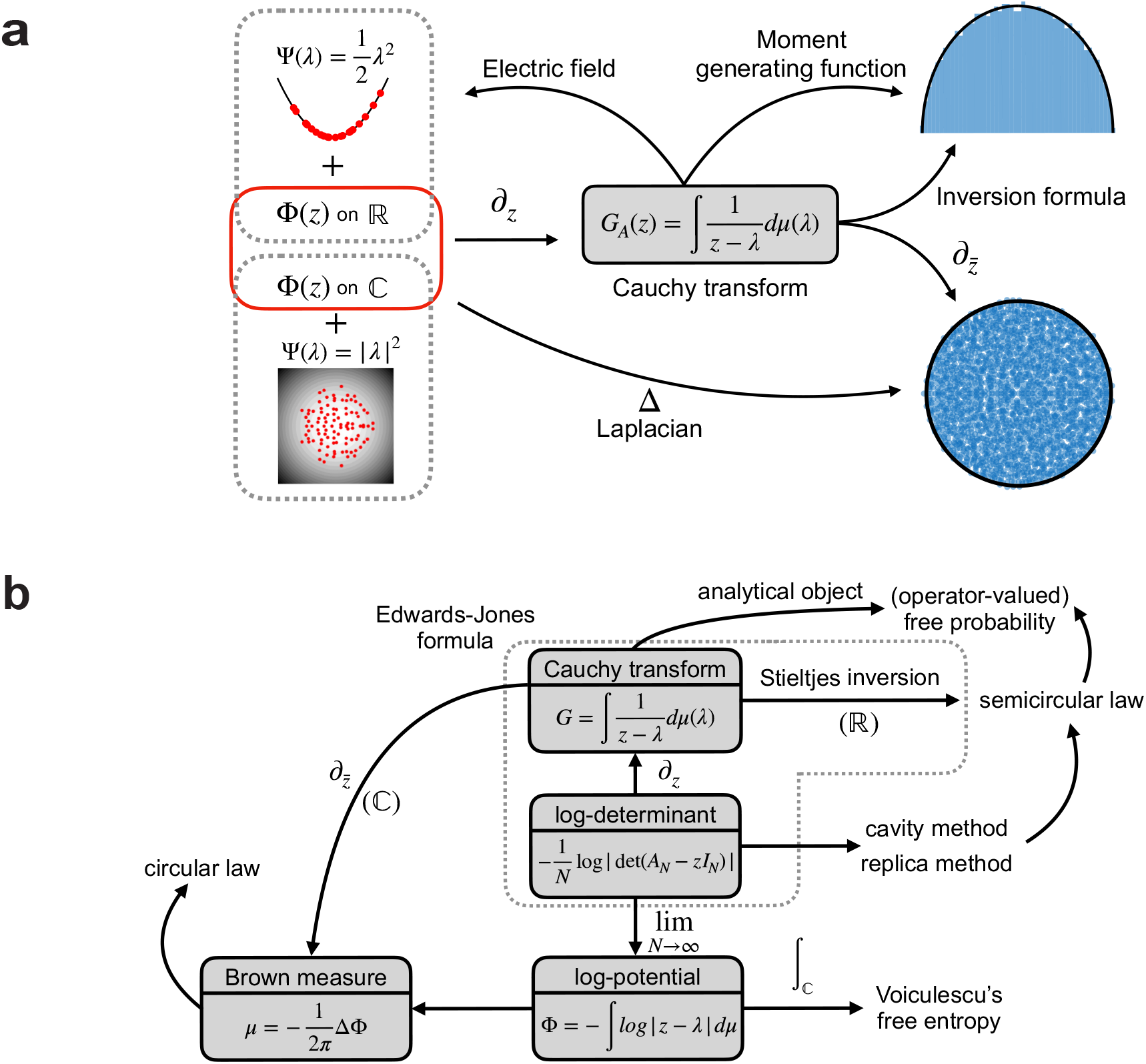
Key mathematical tools used to compute the eigenvalue density *µ*(*λ*). **a)** Concept map depicting the derivation of the semicircular and circular law from the logarithmic potential in ℝ and ℂ. Gray dotted lines – potential terms in the energy functional on ℝ and ℂ. Red solid lines – logarithmic potentials. **b)** Relationships between concepts in random matrix theory (Section IV).

A minimization of total energy involves balancing the forces that confine eigenvalues together and push them away from one another (Figure 3a). As *N* grows to infinity, eigenvalues concentrate at this minimum and converge to a deterministic distribution. This high-dimensional phenomenon, known as the *concentration of measure*, is simply a consequence of when deviations from an average become small with system size. A popular, linear example of this is the law of large numbers. When controlling for variance, a limiting, deterministic distribution exists for the eigenvalues at minimal energy. Our next goal is to find this minimum.

### B. Introducing the Cauchy transform

Finding the minimum energy configuration is possible by differentiating the continuous energy functional *I*[*µ*] with respect to the limiting measure *µ*(*λ*), then differentiating again with respect to *λ*. Many approaches to recovering the density *µ* follow after the second differentiation, but we highlight here that the logarithmic potential Φ(*z*) differentiates to:

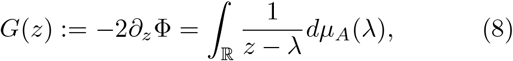

where *∂*_*z*_ is the Wirtinger derivative 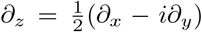 (Figure 3a). *G*(*z*) is defined on ℂ \ ℝ and is called the *Cauchy transform* (also known as the Stieltjes transform, with a sign change, or the Green’s function). The Cauchy transform *G* : ℂ^+^ → ℂ^*−*^, where ℂ^+^ := {*z* ∈ |ℂ ℑ (*z*) *>*0} and ℂ^*−*^ := {*z* ∈ℂ | ℑ (*z*) *<* 0}, cannot be defined on the support of *µ*_*A*_ because of its poles. In the context of Coulomb gases, it is the complex electric field (Figure 3a). If *µ*_*A*_ is compactly supported, meaning that the density is only non-zero on a restricted and finite region *µ*_*A*_([−*r, r*]) = 1 for positive real number *r*, then the Cauchy transform has a power series expansion

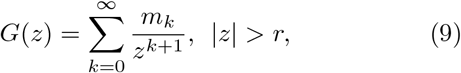

where 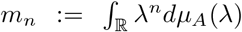 are the moments of *µ*_*A*_. Hence, 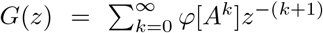 is also a type of moment generating function (Figure 3a). If we take the usual moment generating function to be 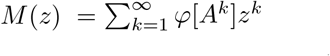, then the relationship between *M* (*z*) and *G*(*z*) is:

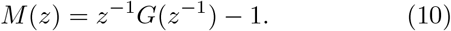

We will use this relationship to convert equations of *G*(*z*) into equations of the moments of *µ*_*A*_ in networks with unstructured variance.

Based on the definition of the linear functional *φ*, the Cauchy transform from Equation (8) can be rewritten as:

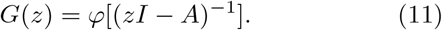

The matrix argument (*zI* − *A*)^−1^ is termed the resolvent of *A*. Thus, random matrix theory tools that involve (*zI* − *A*)^*−*1^ or similar forms are typically called resolvent methods. If one can calculate *G*(*z*) directly in this way, one also circumvents the need for the log-potential. We will be using an expansion of the resolvent to calculate moments in networks with structured variance.

### C. Recovering the spectral density: Hermitian matrices

There are numerous ways to solve for *µ*(*λ*) given *G*(*z*) or Φ(*z*), but most analytic tools fall into two camps. The first group of methods leverages the Sokhotski-Plemelj formula [64, 68]:

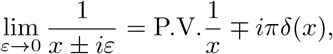

where P.V. is the principal value and *ε* ∈ ℝ_+_. One can easily spot that the left hand side matches the integrand of the Cauchy transform. Application of the formula derives the well-known Stieltjes inversion formula and recovers the spectral density,

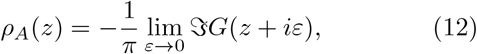

which we will use for the numerical verifications in our work. Another version of this formula, called the Edwards-Jones formula [68, 69], involves the derivative of the log characteristic polynomial, which also evaluates to the Cauchy transform (Figure 3b). Other methods study the eigenvalues as stochastic processes (heuristic method based on Dyson Brownian motion) or recover the density through the moments of *A* (a combinatorial method) [61, 70].

Through any resolvent method, one would find the Cauchy transform of our GOE example is 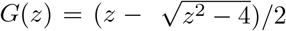. Applying the Stieltjes inversion formula, the density is found to be a centered semicircle on the real line (Figure 3a). The density of the locations of interacting Coulomb charges in the presence of an external potential is in its minimal energy configuration under the semicircle law. Remarkably, this result is true for any Hermitian matrix with i.i.d. entries (apart from symmetry) and finite variance, regardless of the distribution of entries. This is known as the *universality* of Wigner’s semicircular law, covering both local statistics within the bulk as well as fluctuations at the edge of the support under additional conditions. Universality proves that macroscopic behavior in the global density of eigenvalues does not depend on microscopic structures from the specific distribution of matrix entries.

### D. Recovering the spectral density: non-Hermitian matrices

Another centerpiece universality result in random matrix theory is that the normalized limiting eigenvalue distribution of non-Hermitian matrices with i.i.d. entries is a uniform unit disk on ℂ [66, 71–73]. The universality result holds even under certain structured dependencies, such as exchangeability [71] or sparsity [72]. Consider non-Hermitian matrix *B* with i.i.d. real Gaussian entries, known as a real Ginibre ensemble. The support of the spectrum of *B* is no longer limited to the real line, yet the Stieltjes inversion formula relies on discontinuities along ℝ. We can apply *G*(*z*) on ℂ just like we did on ℝ, but we would now need the information of *G*(*z*) inside the support.

Previously mentioned methods fail as soon as random matrices are non-normal [66]. While non-Hermitian matrices may be normal, we will use non-Hermitian matrices as a vehicle to discuss the second camp of analytic tools. Take 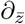to be the conjugate Wirtinger derivative 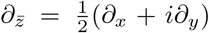. Immediately we can see that 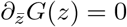 everywhere except at its singularities, meaning the Cauchy transform is holomorphic everywhere except on the support of *µ*. Contributions from the singularities on the support can be found using the fundamental solution for the operator 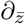, and we have the result

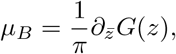

which is also equivalent to Gauss’s law. If we have the Laplacian 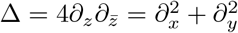, then

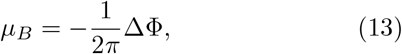

with the log-potential now integrating over ℂ rather than ℝ. Then, one can directly calculate the spectral measure from Φ, without using the Cauchy transform (bottom arrow in Figure 3a). Equation (13) is the definition of a Brown measure, a generalization of the spectral measure for non-normal operators (Figure 3b) [24, 66]. This approach is preferred for non-Hermitian matrices because the log-potential Φ, integrating over its spectral measure, can be rewritten with respect to the measure of its singular values. The relationship between the measures of singular values and eigenvalues is strictly determinantal, so Φ is often calculated as

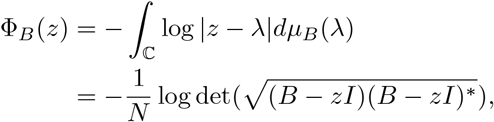

a method known as Girko’s Hermitization (Figure 3b) [24, 25, 66, 74]. In contrast, the Cauchy transform lacks a Hermitization identity to convert the measure of interest to singular values, hence the appeal of the log-potential. Although, one can still apply Hermitization and derive a self-consistent equation of the Cauchy transform (Green’s function in the quantum physics literature) using self-energy functionals, called the Dyson equation [75–77]. In our work, however, we will stick to the Stieltjes inversion formula (Equation 12) because the central object of our study will be a covariance matrix, which is Hermitian.

## V. FREE PROBABILITY

### A. Basic definitions

The central question in free probability asks how to study variables that do not commute, and thus how to define the algebraic structure behind universality. Non-commutative matrices *A* and *B*, for example, render *φ*[*ABAB*] ≠ *φ*[*AABB*]. The joint distribution of *k* non-commutative random variables can no longer be captured by a single commutative measure on ℝ^*k*^, where the distinction between *φ*[*ABAB*] and *φ*[*AABB*] does not exist. We require new mathematical techniques to handle this situation.

First, we define a non-commutative probability space (*C, φ*) to have a unital algebra *C* (with a norm and involution, over ℂ) and a unital linear functional *φ* : *C* →ℂ, with unital meaning *φ*[1] = 1. In the context of random matrix theory, we take *φ* to be the expected normalized trace. For compactly supported measures, such as the semicircle law, what we intuitively do is expand the collection of moments to include all non-commutative forms. Specifically, moments of an element *A* ∈ *C* are (*φ*[*A*^*k*^])_*k∈*N_, and the collection of joint moments of (*A*_1_, …, *A*_*n*_) with *n* ∈ ℕ is the collection of all numbers *φ*(*A*_*σ*(1)_…*A*_*σ*(*k*)_) where *σ*(1), …, *σ*(*k*) ∈ {1, …, *n*}. The collection of joint moments simply consists of every possible product of the powers of *A*_1_, …, *A*_*n*_, and this collection constitutes the (non-commutative) joint distribution of (*A*_1_, …, *A*_*n*_). We must resort to a combinatorial object because it is still unclear as to what a joint ‘non-commutative probability measure’ is [78].

In the classical setting, the joint moments of two independent random variables factor into the product of their marginal moments. The property of *freeness* is independence extended to non-commutative random variables. Excluding some theoretical details pertaining to subalgebras, we say two non-commutative random variables *A* and *B* in (*C, φ*) are *free* if for any polynomials *p*_1_, …, *p*_*n*_ and *q*_1_, …, *q*_*n*_ that satisfy

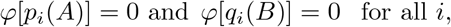

we have that

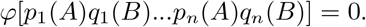

In other words, two random variables are free if the alternating products of centered elements are centered, or if the mixed moments of these centered elements vanish.

As a simple example, say we have two, uncentered free random variables *A, B* ∈ *C*. Then the following are true:

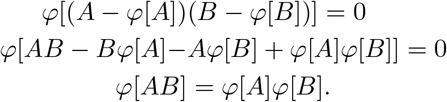

Therefore, freeness enables the calculation of joint moments from marginal moments, as is enjoyed for independent scalar random variables. Here, the factorization is identical to that of independent variables in classical probability, but this is not always true (e.g. *φ*[*ABAB*] = *φ*[*A*^2^]*φ*[*B*]^2^ +*φ*[*A*]^2^*φ*[*B*^2^] −*φ*[*A*]^2^*φ*[*B*]^2^ ≠ *φ*[*A*^2^]*φ*[*B*^2^], see Figure 4a). Another definition of freeness is that their mixed free cumulants vanish – this definition simplifies both assumptions requiring centered and alternating elements. Free cumulants are a structural mirror to classical cumulants, defined on a different set of partitions (non-crossing vs. all set partitions).

**Figure 4.**
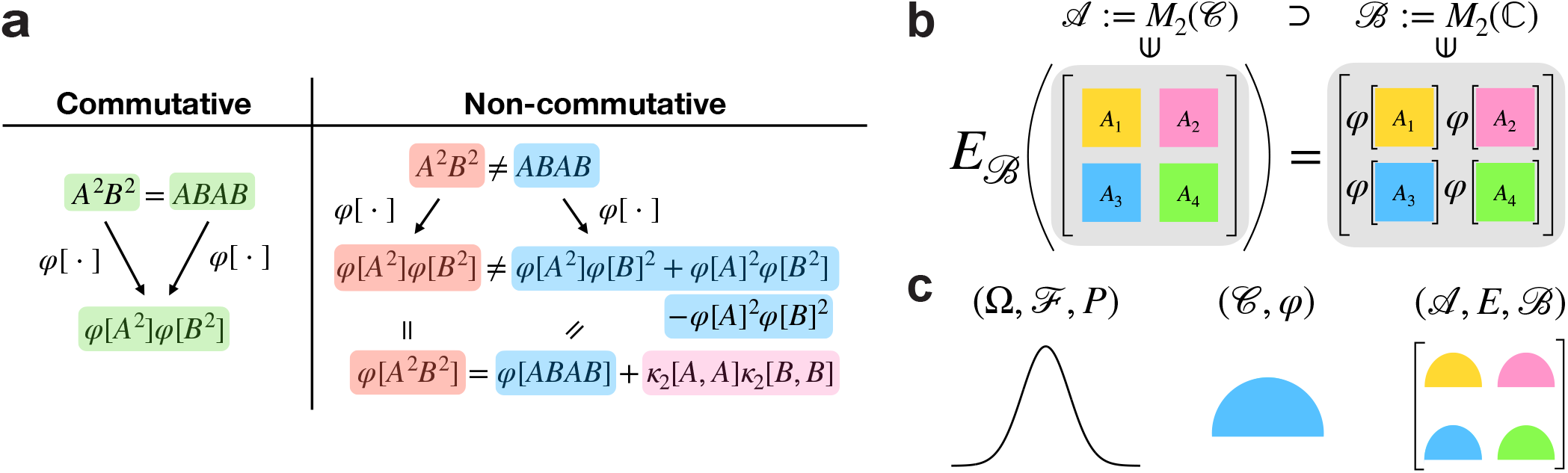
Concepts in free probability theory. **a)** Comparison between expectation calculations of commutative and non-commutative random variables. In the schematic, *κ*_2_[*·,·*] is the multilinear free cumulant functional for two arguments. Since both arguments of *κ*_2_ are the same, this is the second free cumulant, or “variance”, for matrix *A* or *B*. While we can certainly write the pink term in terms of moments instead, following an exact moment-cumulant formula, we use cumulants to avoid redundancy and for concision. **b)** Illustration of the conditional expectation with respect to *B* = *M*_2_(ℂ), operating on elements from *A* = *M*_2_(*C*). **c)** In classical probability, the central element (limiting distribution of the CLT) is the Gaussian variable, defined in probability space (Ω, *F, P*), where Ω is the sample space, *F* is the algebra of events, and *P* is the probability measure. In free probability, the analogue is the semicircular element in the non-commutative probability space (*C, φ*), where *C* is a unital algebra with a norm and involution (e.g. a *C* ^*^*−* algebra) and *φ* is a unital linear functional. In operator-valued free probability, the parallel is the *B−* valued semicircular element in operator-valued non-commutative space (*A, E, B*), where *A* is a *C* ^*^ *−*algebra, *B* ⊗ *A*, and *E* is the conditional expectation defined with respect to *B*. The example shown is a *M*_2_(ℂ)*−*valued semicircular element, where each block is itself a semicircular element in (*C, φ*).

The classical central limit theorem (CLT) states that the normalized sum of i.i.d., centered, commutative random variables with finite variance converges to a normal random variable. Its free probability counterpart, the free CLT, states that the normalized sum of free and identically distributed (f.i.d.), centered, non-commutative random variables with finite variance converges to a semi-circular variable. As such, the semicircle is seen as the free analog of the Gaussian distribution. While in classical probability the normal distribution maximizes differential entropy under second moment constraints, in free probability settings, the semicircle maximizes a relative entropy that contains Voiculescu’s free entropy [24, 79]. Additionally, only the second cumulant of a centered Gaussian distribution is nonzero, and only the second free cumulant of a semicircle distribution is nonzero.

### B. Example: A + B

In both classical and free probability, the mixed cumulants of two independent or free random variables vanish.

We explore this property with the example of the sum *A* + *B*, where *A* and *B* are free random variables. We aim to show that the cumulants of *A* + *B* depend only on the cumulants of *A* and the cumulants of *B*, and to provide a path to computing the Cauchy transform of *A* + *B*: *G*_*A*+*B*_(*z*).

If *A* and *B* are compactly supported, then the moments of *A* + *B* are also uniquely determined by a compactly supported probability measure. The joint distribution of free variables is also determined by their marginal distributions and not on the concrete realizations of the random variables. Letting *K*(*z*) be the inverse function of the Cauchy transform *G*(*z*) such that *G*(*K*(*z*)) = *z*, we define the *R*-transform to be:

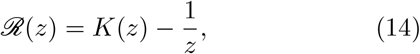

and using moment-cumulant formulas, we can find that

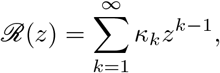

where *κ*_*k*_ is the *k*th free cumulant. While the Cauchy transform is a form of the moment generating function, the *R*-transform is the cumulant generating function. Its application on the Cauchy transform is also equivalent to self-energy in quantum field theory [80, 81]. In the same way that the cumulants of two independent variables add, if the two free variables *A* and *B* with compactly supported measures have corresponding *R*-transforms *R*_*A*_(*z*) and *R*_*B*_(*z*), then the *free additive convolution* of the variables is

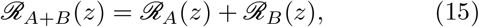

and *R*_*A*+*B*_(*z*) gives the *R*-transform of the sum. Thus, free cumulants linearize convolutions in both settings.

Using Equations (14) and (15), we next show how one can obtain the limiting spectral distribution of *A* + *B* utilizing the *R*-transforms and Cauchy transforms of *A* and *B* individually. Letting the Cauchy transform of *A* be *G*_*A*_(*z*) and the inverse function of *G*_*A*_(*z*) be *K*_*A*_(*z*), a simple calculation gives:

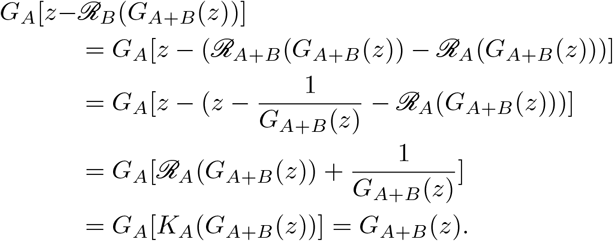

The relationship

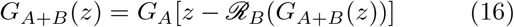

is the *subordination property*. If *G*_*A*_ and *R*_*B*_ are known, this self-consistent equation can be solved numerically for *G*_*A*+*B*_(*z*), followed by an application of the Stieltjes inversion formula. Special cases in this setting contain explicit solutions if one can identify the correct solution out of the multiple given by nonlinearities, but a general solution eludes us. However, a nontrivial explicit solution for *G*_*A*+*B*_(*z*) in the operator-valued setting, introduced in the next section, is an extreme rarity [24]. Nevertheless we can find the first few moments of *G*_*A*+*B*_(*z*) analytically (as we will show in later sections).

## IV. OPERATOR-VALUED FREE PROBABILITY

So far, we have mentioned Hermitian and non-Hermitian examples of random matrices (Gaussian and Ginibre ensembles) with foundational limiting distributions. We also discussed that their universality holds even under specific dependencies [71, 72]. Another type of dependency between entries is block structure, where each block may be its own ensemble and blocks may be repeated. Unfortunately, the free probability tools introduced in the previous section cannot properly handle block matrices, particularly with blocks in repeated positions. Intuitively, calculating the moments of a block matrix would involve the bookkeeping of both entry-level interactions and block residence. In addition, collapsing the information of the matrix to a scalar through the expected normalized trace *φ* ignores any information about its block structure. For example, even if a Hermitian block matrix has free semicircular elements for its blocks, the whole matrix may not be semicircular with respect to *φ*. A new theoretical framework must be introduced for such elements.

The natural solution to this problem is to stop insisting on a scalar answer at each calculation, even if the end goal is to find the scalar eigenvalue distribution of the block matrix. Preserving the block structure in determining the moments of block matrices is an application of *operator-valued free probability*, where block matrices are one example of an *operator-valued element*. Let ⊗ be the Kronecker product and **A** ∈ *A* = *M*_*d*_(*C*) : *M*_*d*_(ℂ)⊗ *C* be a *d* × *d* block matrix with each block being its own matrix from *C*. Instead of applying *φ* on the whole matrix **A**, we apply *φ* to each block of **A**:

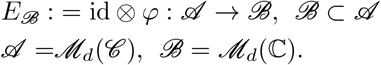

This block-level operation *E*_*B*_ (or *E*, dropping the subscript for brevity) is called the *conditional expectation*, and is defined with respect to a subalgebra *B* (Figure 4b). Compared to the free probability setting, ℂ is replaced with a non-commutative subalgebra *B*, and *φ* is replaced by *E*_*B*_. Rather than a scalar-valued non-commutative probability space (*C, φ*), we now construct an operator-valued non-commutative probability space (*A, E, B*) (Figure 4c).

If moments are defined with the conditional expectation in operator-valued free probability, similar to how *φ* defines moments in free probability, it is transparent that now, moments are no longer scalar quantities and thus no longer commute. Crucially, this change implies that, for **A, B** ∈ *A, E*[**A**] ≠ *E*[**B**] *E*[**B**]*E*[**A**]. For multiplication by elements in *B* (equivalent to scaling by elements from ℂ), we would have the bimodule property *E*[*B*_1_**A***B*_2_] = *B*_1_*E*[**A**]*B*_2_, for *B*_1_, *B*_2_ ∈ *B*. Also, *E*[*B*_1_] = *B*_1_, similar to how E[*c*] = *c* for *c* ∈ ℂ. On the other hand, *B*_1_ must remain nested in *E*[**A***B*_1_**A**]. This complicates the definition of a distribution, which we defined in the previous section as the collection of moments. Instead of the collection (*φ*[*A*^*k*^]) for *A* ∈ *C*, one needs to consider all *operator-valued moments* of **A**:

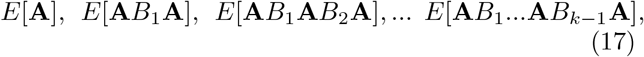

for *B*_1_, …*B*_*k−*1_ ∈ *B*. This collection constitutes the *operator-valued distribution* of **A**.

Taking into account the new conditional expectation and structure of operator-valued moments, there are few modifications to be made for many of the definitions in the previous sections. Freeness, for instance, is still a question of whether the alternating products of centered polynomials of two operator-valued random variables are also centered. Only, the polynomials have coefficients from *B* and the powers contain sandwiched elements from *B*, like those in Expression (17). The alternative definition, where the vanishing of mixed operator-valued cumulants implies freeness, is also the same in this context. To disambiguate freeness in the two settings, we say two operator-valued variables are *free with respect to E*_*B*_ instead, as the polynomials are defined according to *B*. While the crux of the definition is unchanged, the dependence on *B* implies freeness may not persist across subalgebras.

The simplest illustration compares an arbitrary subalgebra *B* ⊗ *A* with the minimal subalgebra ℂ ⊗ *B* ⊗ *A*. *A* is technically both a scalar-valued and an operator-valued probability space, depending on whether the space is defined with respect to tr ⊗ *φ* : *A* → ℂ or *E*_*B*_, respectively. Due to the difference in the commutativity of moments in the two probability spaces, two block matrices in *A* can be free with respect to *E*_*B*_, but not with respect to tr ⊗ *φ*. Therefore, computations may be less tractable in ℂ, which is ultimately a reduction to the scalar-valued free probability setting, than in *B*, the larger subalgebra.

For symmetric block matrix **A**, we define its operator-valued Cauchy transform *G*_**A**_ : ℍ^+^(*B*) → ℍ^*−*^(*B*) to be

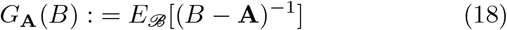

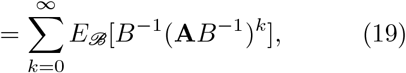

where ℍ^+^(*B*) := {*B* ∈ *B* |ℑ *B*) *>* 0}and ℍ^*−*^(*B*) := {*B* ∈ *B*|ℑ (*B*) *<* 0 are the operator upper and lower half-planes. The structure of this definition follows the scalar version (Equation 11), only the domain changes.

Equation (19) shows that *G*_**A**_ acts as a moment generating function of **A**. Similarly, the relationship between the Cauchy transform and the *R*-transform holds (Equation 14), and thus so does the subordination property (Equation 16). The operator-valued *R*-transform is still the cumulant generating function, only the terms are operator-valued cumulants with respect to *B*.

Equation (19) shows that *G*_**A**_ acts as a moment generating function of **A**. Similarly, the relationship between the Cauchy transform and the *R*-transform holds (Equation 14), and thus so does the subordination property (Equation 16). The operator-valued *R*-transform is still the cumulant generating function, only the terms are operator-valued cumulants with respect to *B*.

Does the benchmark distribution of free semicirculars in scalar-valued free probability also have an operator-valued counterpart? The analogous operator-valued semicircle distribution arises concretely as the limiting distribution of the operator-valued free CLT. The normalized sum of centered f.i.d. Hermitian operator-valued variables converges to a distribution that has moments and self-consistent Cauchy transform equations comparable to those of a semicircular element. These moments must depend on *B*, and so this distribution is termed a *B*−valued semicircular distribution, and an element with such a distribution is called a *B*−*valued semicircular element*. An element is *B*−valued semicircular if all operator-valued cumulants except the second are zero, identical to the semicircular variables in scalar free probability and Gaussian variables in classical probability.

The spread or second moment of the limiting distribution of the operator-valued free CLT can no longer be captured by a scalar, meaning we must also revisit how variance is portrayed in the operator-valued setting. We define the covariance mapping *η* : *B* → *B* as:

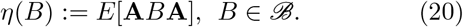

Notice how if *B* = ℂ, *η*(*c*) = *cφ*[**A**^2^] for *c* ∈ ℂ, and the mapping only depends on the second scalar moment of **A**. If *B* ≠ ℂ, the *B* valued semicircle distribution cannot be fully characterized by a single number, but only by how a variable with such a distribution interacts with all elements of *B*. Then, since all higher cumulants vanish, the *R*-transform of a *B*−valued semicircular element *R*_**A**_(*B*) is simply the covariance map: *R*_**A**_(*B*) = *η*(*B*).

Each transition, from classical probability to free probability, then from scalar-valued to operator-valued free probability, involves an additional layer of non-commutative objects. The consequences of added non-commutativity, luckily, produce rather few changes in the applications of the analytical objects in each setting. The CLT, linearization through cumulants, and notions of independence, are all shared (Table I). Rather, it is the spaces in which these objects lie that add complication. In Section III, we laid the groundwork for a mechanistic circuit with random connectivity. As we will see in the following section, determining the statistics from this circuit presents a very natural environment to use (operator-valued) free probability.

**Table I.** Comparison table of concepts in classical probability, free probability, and operator-valued free probability.

| Classical probability | Free probability | Operator-valued free probability |
| --- | --- | --- |
| Everything commutes | Moments commute<br>variables don't | Neither variables nor<br>moments commute |
| Independence | Freeness | Freeness wrt<br>conditional expectation |
| Gaussian | Semicircular | $\mathcal{B}$-valued<br>semicircular |
| CLT | Free CLT | Operator-valued<br>free CLT |
| Convolution | Free convolution<br>with $\mathcal{R}$-transform | Operator-valued<br>$\mathcal{R}$-transform |

## VII. RANDOM NETWORKS WITH NO GLOBAL STRUCTURE

Now armed with new free probability techniques we return to our recurrent network introduced in Section Equation (5) provides an explicit calculation of our object of interest, *C*, in terms of random matrices *R* and *W* :

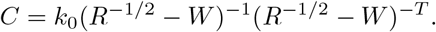

In this section we aim to calculate the eigenvalue density *µ*(*λ*) of *C*, and provide a compact relation between the effective dimension *D*(*C*), the uniformity of firing rates *D*(*R*), and the variance of synaptic connections *g*^2^*/N*.

### A. Free independence of *R* and *W*

Application of our free probability tools on Equation (5) in the unstructured network requires the assumption that *R* and *W* are free. In networks with global structure or connectivity matrices with block structure, *R* must be free from the blocks of *W*. In this brief subsection, we discuss the implications of this assumption.

In defining the relation *r*_0_ = *f* (*µ* + *Wr*_0_), we have demonstrated that *R* depends on *W*, and thus they cannot be independent of one another. To account for the influence of local interactions within the circuit, one would typically solve for *r*_0_ self-consistently through this equation. In our work, however, we will make the strong statement that *r*_0_ = *f* (*µ*). In other words, we assume that firing rates are purely driven by external inputs, that the elements of *W* are weak and have little influence on *r*_0_. In support of this statement, the stimulus tuning of neurons in the sensory cortex is primarily established by feedforward pathways which carry information that is external to the network [82]. However, if *W* is considered negligible, Equation (5) would be reduced to *C* = *k*_0_*R*, and neuronal activity is uncorrelated (i.e. *C* is diagonal). To avoid this consequence, we will assume that while *R* is not determined by *W, C* is very dependent on *W*. This amounts to taking recurrent interactions as the sole source of pairwise covariability, as is often done in modeling studies [83].

These arguments, while purporting an idealized view of recurrent circuit dynamics when driven by external inputs, are needed so that we can properly handle *C* as a function of free matrices *R* and *W*.

### B. Stability of *C*

Taking *R* and *W* as free allows us to determine the stable regime of *C*. For the steady-state covariance matrix to exist, we would need the series

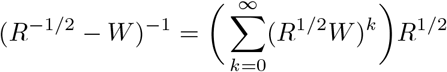

to converge. The operating regime of the system is where the spectral radius *ρ*(*R*^1/2^*W*) *<* 1. Now, recall that *W* is an *N* × *N* matrix with i.i.d. real Gaussian entries, or a realization of a scaled, real Ginibre ensemble. As discussed in Section IV, the distribution of eigenvalues in the large *N* limit for a Ginibre ensemble is a uniform disk on the complex plane, so *W* in this limit is a circular element. A real Ginibre ensemble *W* is biorthogonally invariant, meaning if *U* and *V* are orthogonal matrices then *UWV* and *W* are equally distributed. A biorthogonally invariant matrix has a measure satisfying the single ring theorem, which finds that in the large *N* limit, the measure converges to an annulus centered at the origin in ℂ (this contains the circular law). The single ring theorem also states that the outer radius of the annulus is given by the density of singular values [81, 84, 85]:

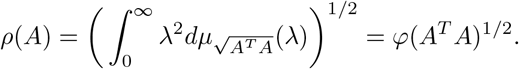

While the product *R*^1/2^*W* is only right orthogonally invariant, its eigenvalues have the same distribution as that of a biorthogonally invariant matrix. Therefore,

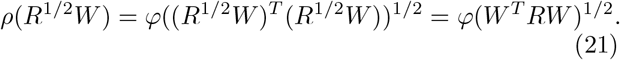

Since *R* and *W* are asymptotically free and the trace operation is cyclic, we have

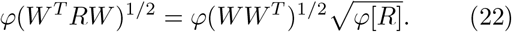

The matrix *WW*^*T*^ is a scaled, real Wishart ensemble. It has a limiting distribution equivalent to the Marchenko-Pastur (in RMT terms) or free Poisson (in free probability terms) distribution of order 1 [24, 63, 64]. All cumulants of a free Poisson distribution of order *c* (ratio between rows and columns) are equal to *c* [24], so 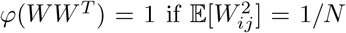 and *W* is square. Since our matrix is scaled,

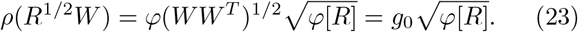

Therefore, the stability condition for the network activity in an unstructured network is simply *g*_0_ *φ*[*R*] *<* 1. If *W* is a 2 × 2 block matrix:

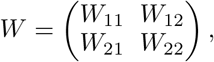

where each block *W*_*αβ*_ (*α, β* 1, 2) is its own ensemble with entries drawn from 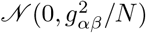, the stability condition changes to

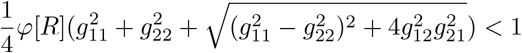

if we first condition on a fixed realization of *R* [86, 87]. From this point, we shall only use variance profiles in the convergent regime.

### C. Eigenvalue density of the covariance matrix

The effective dimension *D*(*C*) is a dimensionless property, so setting *k*_0_ = 1 will not alter *D*(*C*). To determine the eigenvalue density of *C* = (*R*^−1/2^ − *W*)^−1^(*R*^−1/2^ − *W*)^*−T*^, we consider the non-commutative probability space (*C, φ*) where *R* and *W* are free variables. Note that technically *R* and *W* are matrices of finite size in the way we introduced them in the previous section, but we shall use the same notation for their large *N* limit counterparts in *C* to avoid confusion. To fully leverage their freeness, we first construct an operator-valued probability space (*A, E, B*) where *A* := *M*_2_(*C*), *B* := *M*_2_(ℂ) ⊂ *M*_2_(*C*), *E* := *id* ⊗ *φ* : *M*_2_(*C*) → *M*_2_(ℂ). In *A*, we define the symmetric 2 × 2 block matrices **R^−1/2^** and **W** ∈ *A* := *M*_2_(*C*):

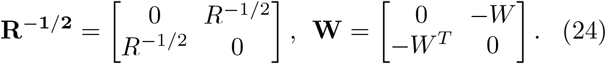

Letting *T* = *R*^−1/2^ − *W* and **T** = **R**^−1/2^ + **W**,

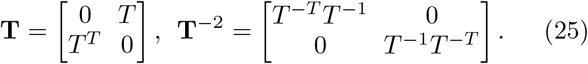

This construction reflects two ideas: Hermitization (*W* is not Hermitian, but **W** is) and deterministic equivalence, whereby functions of random ensembles converge, for large *N*, to the same functions applied to their corresponding limiting operators (Chapters 9–10 in [24]). Finding the expected normalized trace along the diagonal block matrix **T**^−2^ to obtain the moments of its spectral density is equivalent to applying *φ* to either of the diagonal blocks because the nonzero eigenvalues of *T* ^*−T*^*T*^−1^ and *T* ^−1^*T* ^*−T*^ are equivalent. Therefore, the eigenvalue distributions of *C* and **T**^−2^ are identical, and their Cauchy transforms *G*_*C*_(*z*) and 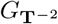 (*z*) are related by the equation

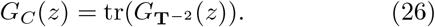

The normalized trace must be taken because the Cauchy transform of **T**^−2^ is operator-valued, lying in *B*, while the Cauchy transform of *C* is a scalar complex function in ℂ. This construction enables us to avoid dealing with the non-symmetric operator *W* and instead work with symmetric operators, where the subordination property for the operator-valued Cauchy transform is well-developed. Additionally, given that the entries of *W* are Gaussian, *W* and −*W* have the same eigenvalue distribution. We will consider **W** with off-diagonal blocks consisting of *W* and *W*^*T*^ rather than −*W* and −*W*^*T*^ for convenience.

Recall that the covariance function *η* : *B* → *B* is defined as *η*(*B*) = *E*_*B*_(**A***B***A**) with **A** being a *B* valued element and *B* ∈ *B*. From [24], we have the fact that **W** is semicircular with respect to *B* because **W** is self-adjoint and the off-diagonal elements *W* and *W*^*T*^ are circular. Since **W** is *B*−valued semicircular, the *R*-transform of **W**, *R*_**W**_(*B*), is equal to *η*(*B*). Furthermore, operator-valued semicircularity allows us to find a smaller subalgebra *D* within *B* such that **W** is also *D* –valued semicircular. We will take *D* = *D*_2_(ℂ) ⊂ *M*_2_(ℂ) to be the subalgebra of 2 × 2 diagonal block matrices. If *η*(*D*_2_(ℂ)) ⊂ *D*_2_(ℂ), then a *B*−valued semicircular element is also a *D*_2_(ℂ)−valued semicircular element, restricting both the covariance mapping *η* and conditional expectation *E* to *D*_2_(ℂ). In other words, if applying the covariance function *η* on diagonal block matrices outputs diagonal block matrices, we can restrict all our calculations to only diagonal block matrices, which are much easier to handle.

This motivation prompts the following calculation:

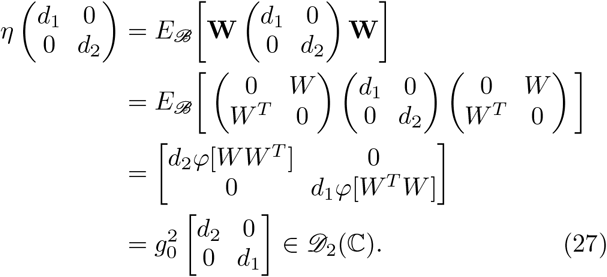

Consequently, **W** is a *D*_2_(ℂ)−valued semicircular element, *E* : *A* → *D*_2_(ℂ), and *η* : *D*_2_(ℂ) → *D*_2_(ℂ). The restriction of the conditional expectation,

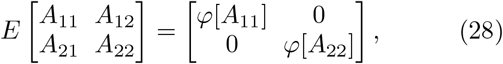

implies that 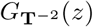 is a 2 × 2 diagonal matrix, as the conditional expectation defines the operator-valued Cauchy transform. From [19], we have the relationship

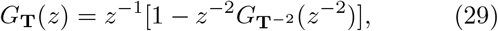

which means *G*_**T**_(*z*) is also diagonal. Let us denote each diagonal component of the Cauchy transforms as below:

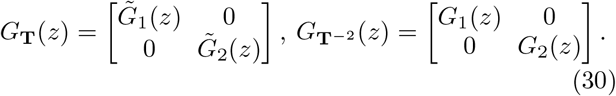

**R**^−1/2^ and **W** are free because their elements *R*^−1/2^ and *W* are free. Then, utilizing the aforementioned subordination property [24] yields:

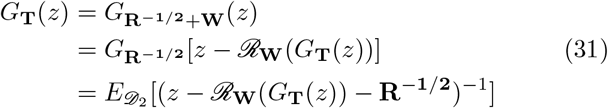

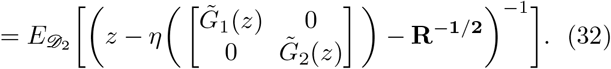

Simplifying the expression inside the conditional expectation and applying the normalized trace results in a system of two equations:

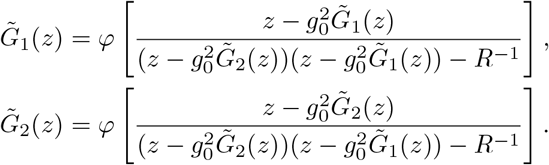

It is easy to see that these two equations are functionally the same, meaning 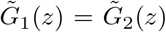, and *G*_1_(*z*) = *G*_2_(*z*). This same result also follows immediately from the fact that *T*^*T*^ *T* and *TT*^*T*^ have the same eigenvalues. Therefore, by Equation (26), 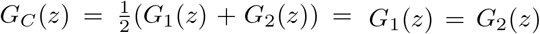. Writing 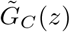, the Cauchy transform is related to *G*_*C*_(*z*) in the same way 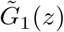 is to *G*_1_(*z*). Then, we have

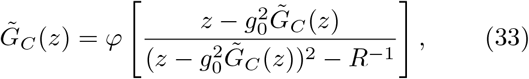

a self-consistent equation for 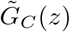, from which the Cauchy transform of *C* follows using Equation (29). Note that *φ* is averaging over randomness originating from *R*, since 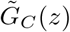 is a scalar. From Equation (33), *G*_*C*_(*z*), and thus the density by the Stieltjes inversion formula, is calculated numerically (see Appendix A for numerical simulations, example shown in Figure 5a).

**Figure 5.**
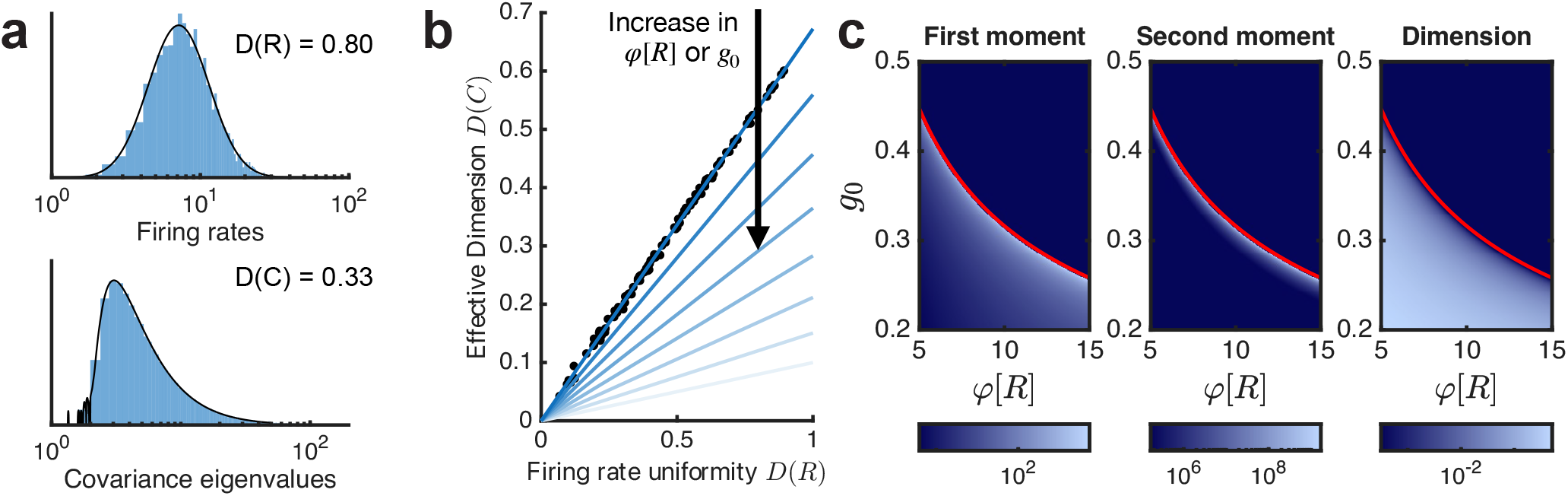
**a)** Example distributions of firing rates {*r*_*i*_} and eigenvalues *λ*_*i*_ for a network with *g*_0_ = 0.1897. The rates were sampled from *LN*(*µ, σ*^2^), with *µ* = 2.191, *σ*^2^ = 0.2231, while the eigenvalue density *µ*(*λ*) (black line) was computed from Equation (33). The associated *D*(*R*) and *D*(*C*) values are given. **b)** The main result from Section VII: *D*(*C*) and *D*(*R*) are linearly related according to 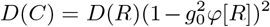 (Equation 36). The larger the magnitude of *g*_0_ or *φ*[*R*], the smaller the magnitude of the slope relating *D*(*C*) and *D*(*R*). Solid lines represent the theoretical result, with lighter colors for larger values of *φ*[*R*] or *g*_0_. Black dots represent simulation results, showing good agreement. The theoretical relationship between *D*(*R*) and *D*(*C*) was plotted using *g*_0_ = 0.1897, *D*(*R*) values between 0 and 1, and *φ*[*R*] from 5 to 19. Simulations were performed with *D*(*R*) sampled uniformly between 0.1 and 0.9, a fixed connectivity matrix *W* with the same *g*_0_, *φ*[*R*] = 5, and 100 samples. **c)** Heatmaps of the first moment *φ*[*C*], second moment *φ*[*C*^2^], and dimension *D*(*C*) of the eigenvalue distribution of *C* across parameter sweeps on *φ*[*R*] and *g*_0_. Solid red lines denote the stability condition 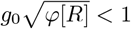.

### D. Calculating the effective dimension of covariability *D*(*C*)

As previously mentioned (Equation 10), the Cauchy transform is related to the moment generating function *M*_*C*_(*z*) by

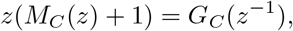

and some quick algebra with Equation (29) gives us the relationship with 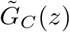:

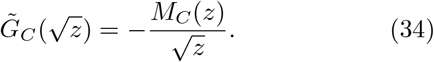

Substituting that expression into Equation (33), we have

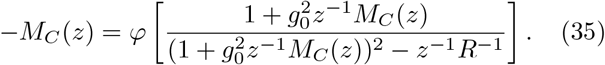

Differentiating this equation once with respect to *z* and setting *z* = 0 gives the first moment of *C*:

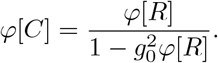

Differentiating twice gives the second moment:

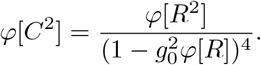

Together, the effective dimension is

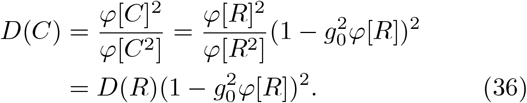

Equation (36) was the central theoretical result in our past work [23]. Specifically, the effective dimension of covariability *D*(*C*) in a neuronal network whose random coupling lacks any global structure depends linearly on the uniformity of the network’s firing rates *D*(*R*). Increasing either the average firing rate *φ*[*R*] or the connectivity variance *g*_0_ decreases the effective dimensionality *D*(*C*) (Figure 5c). We note that if *R* = *I* so that *φ*[*R*] = *D*(*R*) = 1, our result agrees with past calculations [19, 26] where 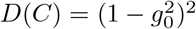.

## VIII. NETWORKS WITH GLOBAL VARIANCE STRUCTURE

For two random matrices to be asymptotically free, it is sufficient if they are independent and one of them is rotationally invariant. The second condition can break down as soon as *W* has structure. For example, neurons in functional modules that process specific types of information are often more likely to be connected [33, 88–92], spatial organization may arise from connectivity probabilities decaying with distance [93–96], or *W* might be a global description of multiple regions of neurons that have different connection properties [29, 30, 97, 98]. In the following sections, we will explore one version of the third scenario, in which the network contains two areas with connections that differ in their variance. We will keep Gaussianity in the entries of *W*, but let the connectivity of each area be independent Ginibre ensembles, as described in Section VII B.

With 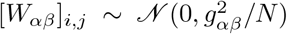, global structures in *g*_*αβ*_ can be constructed in multiple ways: we can vary *g*_*αβ*_ within a single area (*W*_11_ or *W*_22_), vary *g*_*αβ*_ across the two areas (*W*_12_ or *W*_21_), or vary *g*_*αβ*_ across multiple blocks of *W*. Although *R* and *W* are no longer free (i.e. mixed centered moments do not vanish), proper selection of the subalgebra *B*, within which we define the conditional expectation *E*, renders this fact irrelevant. Since each *W*_*αβ*_ is a Ginibre ensemble assumed to be free from *R, W* and *R* are free with respect to the subalgebra *M*_2_(ℂ) [24]. With operator-valued freeness assumed, we will be able to characterize how global structure may change in the theoretical predictions of *D*(*C*).

### A. Added within-area connectivity variance

Without loss of generality, we increase the variance of connections within area 2 by *κ*^2^ (Figure 6a). We will keep the total amount of variance in *W* fixed for all types of added structure, meaning the second moment of the singular value distribution of *W* remains fixed as *κ* varies 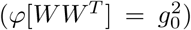. Letting 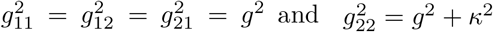, the condition for conserved variance is:

**Figure 6.**
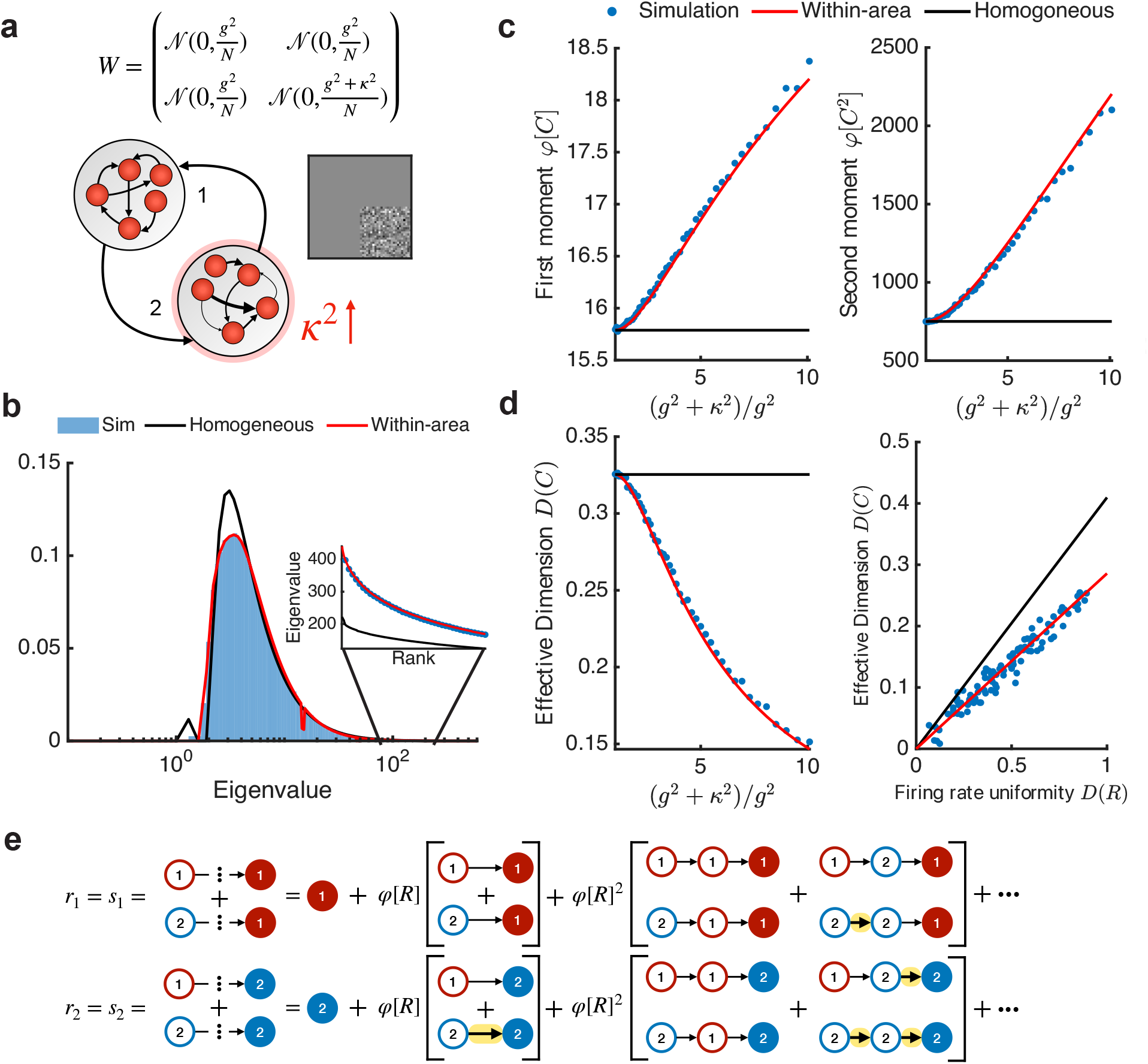
**a)** A recurrent network with two areas and increased variability within area 2. The gray square shows the *κ*^2^ contribution to the bottom right block of the connectivity matrix *W*. *W* becomes a 2 ×2 block matrix, where each block has i.i.d. Gaussian entries. Total variance in *W* is conserved to isolate the effects of added structure, so *g*^2^ decreases as *κ*^2^ increases. **b)** Simulated eigenvalue distribution of covariance matrix *C* against theoretical predictions from Section VII with uniform variance in *W* and Section VIII A with areal-level structure. Theoretical predictions are calculated by solving the self-consistent Cauchy transform equation (Equation 33 vs Equations 39-40) numerically and applying the Stieltjes inversion formula. Solid black line – unstructured connectivity (*κ*^2^ = 0). Solid red line – structured connectivity (*κ*^2^ = 0.07). Inset – largest eigenvalues in the distribution ordered by rank. Blue dots – simulated eigenvalues. Spectral density shows agreement with simulations at both the bulk and the edge. **c)** Theory and simulations of the first and second moments with increasing *κ*^2^. **d)** Left: Theory and simulations for *D*(*C*) across *κ*^2^ match as well. Right: The linear relationship between *D*(*R*) and *D*(*C*) is preserved but the slope is reduced (*κ*^2^ = 0.07). **e)** Components of the sending (*s*) and receiving (*r*) vectors broken down by the length of propagation pathways and the target area. For components *r*_1_ and *s*_1_, for example, we consider all pathways that end in area 1. A factor of *φ*[*R*] is multiplied for each increase in path length. Solid thin arrows represent how 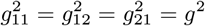, while bold and highlighted arrows represent 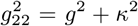. The increase in 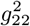 contributes mainly to *r*_2_ and *s*_2_ components, indicating increased alignment between strong receiver and strong sender.

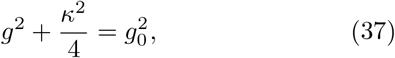

where 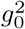 is fixed and *g*^2^ decreases with increasing *κ*^2^. This constraint will allow us to compare across networks with varying degrees of structure (in particular to networks with *κ* = 0). As before, we want to construct **T** such that manipulation of **T** gives the distribution of *C*. Now that *W* is also a block matrix, we set

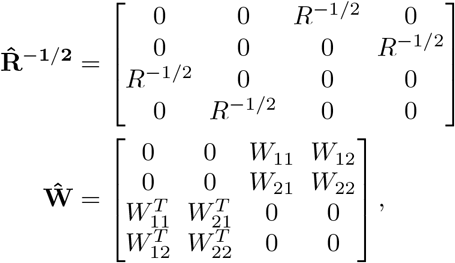

and so 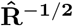 and **Ŵ** are now symmetric 4 × 4 block matrices. Note that even though 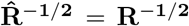, we must make this distinction here such that 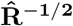 lies in the same algebra as **Ŵ**. The *R*^−1/2^ in Equation (24) is double the size of the *R*^−1/2^ here in the 4 × 4 case, but their variables in the large *N* limit are equivalent.

Accordingly, we define a new operator-valued probability space (*A*′, *E*′, *B*′) where *A*′:= *M*_4_(*C*), *B*′ := *M*_4_(ℂ) ⊂ *M*_4_(*C*), and *E*′ := *id* ⊗ *φ* : *M*_4_(*C*) → *M*_4_(ℂ). Then, 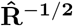, **Ŵ** ∈ *A*′, and we will let 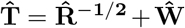. As before, since **Ŵ** is symmetric and its off-diagonals *W*_*αβ*_ are circular, **Ŵ** is *M*_4_(ℂ)−valued semicircular. Therefore, despite the elevation to 4 × 4 block matrices, checking that the covariance mapping 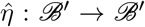 satisfies 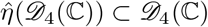 enables us to restrict 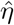 to the subalgebra of diagonal 4 × 4 block matrices in *D*_4_(ℂ):

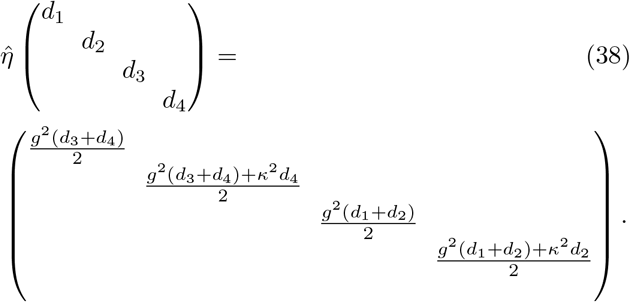

Again, by the definition of an operator-valued Cauchy transform, we must have both *G*_**T**_(*z*) and 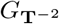 (*z*) ∈ *D*_4_(ℂ). To avoid confusion with notations that have both tildes and hats (for the structured cases), we shall continue the same Cauchy transform notation. Along the diagonals of *G*_**T**_(*z*) and 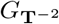 (*z*), we now have 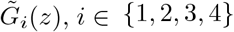 and *G*_*i*_(*z*), *i* ∈ {1, 2, 3, 4}, respectively.

Seeing that *W*_11_, *W*_12_, and *W*_21_ are identical in distribution and thus 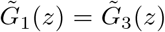 and 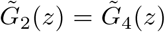, applying the subordination property and simplifying gives a system of two equations

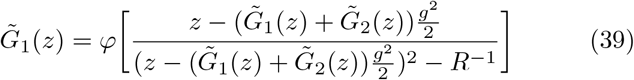

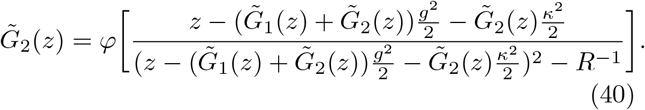

Recall that using the subordination property here is still valid despite block structure in *W* due to the fact that **Ŵ** and 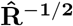 are free with respect to the subalgebra *M*_4_(ℂ). The relationships between *G*_*C*_(*z*) and 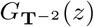 (*z*) (Equation 26) and between 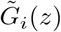 and *G*_*i*_(*z*) (Equation 29) remain the same. Numerical verification of 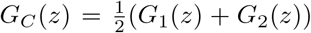 (see Appendix A for numerical simulations) shows excellent agreement between theory and simulations (Figure 6b). In particular, we see that the within-area structure increases the magnitude of large eigenvalues in *µ*(*λ*) (zoom in inset in Figure 6b). We next explore this observation through computing the effective dimension *D*(*C*).

As shown in Equation (25), we can see that the bottom right block of **T**^−2^ is *C*, and subsequently that the bottom right block of **T**^−4^ is *C*^2^. The expansion of the resolvent around *z* = 0 is:

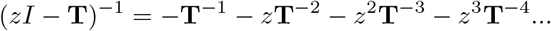

We can immediately see that only even powers contribute to the diagonal blocks of the resolvent, and that these contributions in the lower right block are powers of *C*:

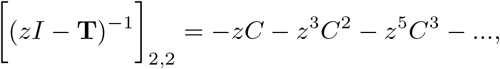

which means

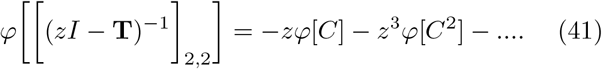

Equation (41) is also an average of the bottom right entries of *G*_**T**_(*z*) or Equations (39)-(40). Expanding the latter allows us to match the *z* and *z*^3^ terms in both Equation (41) and Equations (39)-(40) (details in Appendix B), giving the first and second moments of the eigenvalue distribution of *C*:

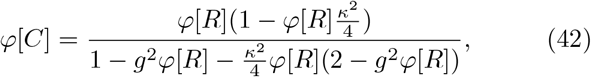

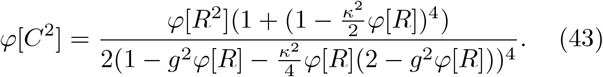

Calculating the participation ratio, the effective dimension is:

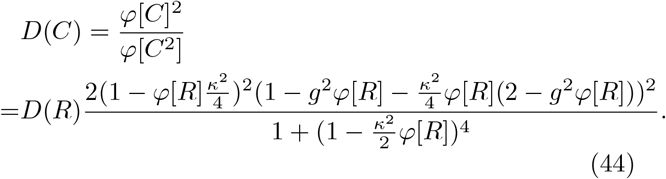

The relation between *D*(*C*) and *D*(*R*) is exact in terms of *κ*^2^ despite *W* and *R* no longer being free. Recall that we only rely on the freeness between **Ŵ** and 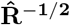 with respect to *M*_4_(ℂ). There is an excellent agreement between the theoretical results in Equations (42), (43), and (44) and numerical evaluation of *φ*[*C*], *φ*[*C*^2^], and *D*(*C*) over a large range of *κ* (Figure 6c,d). Increasing the variance in connectivity within an area increases the first and second moments of the spectral density, but overall decreases the dimensionality. However, the linear relationship between *D*(*C*) and *D*(*R*) is preserved, with the slope decreasing with increasing *κ* (Figure 6d).

Finally, to better compare the relation for within-area structure to the homogeneous case (*κ* = 0) we take an expansion of the denominator in Equation (44) and collect terms to give:

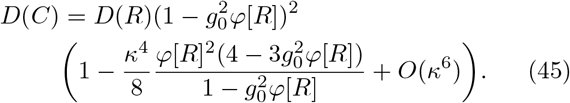

We immediately see how the added variance *κ >* 0 acts to reduce dimensionality compared to the homogeneous case.

To mechanistically understand how a redistribution of connectivity variance controls *D*(*C*), we give attention to the vectors (Appendix B):

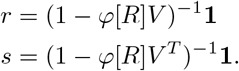

Here, the matrix 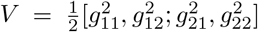 captures the connectivity variance structure. We call *r* and *s* the *receiving* and the *sending* vectors, respectively, because their components demonstrate the variance contributions to and from each area. For example, expanding

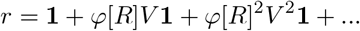

shows the accumulation of variance propagation pathways with increasing length, and ultimately measures variance in response to coupling *V* and gain *R* (Figure 6e). In the within-area case with connectivity variance 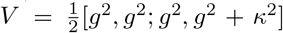, we have that receiving and sending vectors are both equal:

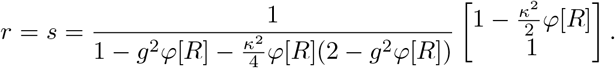

It becomes clear that area 1 (top entry) both receives and sends less variance due to the added variance into the recurrent connectivity in area 2.

The general expressions for *φ*[*C*] and *φ*[*C*^2^] in terms of the components of *r* = [*r*_1_, *r*_2_] and *s* = [*s*_1_, *s*_2_] are (see Equations (B5)-(B6)):

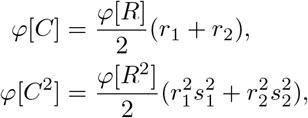

so that we have:

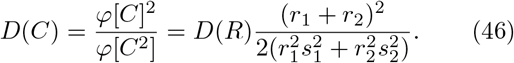

For the within-area case we have the alignment of the sending and receiving vectors, i.e. *s*_1_ = *r*_1_ and *s*_2_ = *r*_2_. This alignment causes 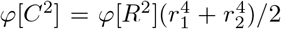. Since *r*_2_ *> r*_1_, the 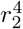 term dominates and *φ*[*C*^2^] increases by a larger fraction than *φ*[*C*]^2^ (Figure 6c), suggesting that a disproportionate amount of variance is concentrated in a few modes.

### B. Added between-area connectivity variance

Now let us consider the case where between-area connectivity *W*_12_ has additional variance (Figure 7a). The connectivity matrix has 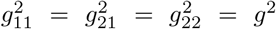 and 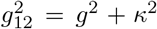, with total variance conserved in a similar way to the previous section. Utilizing the same techniques, the equations for 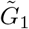 and 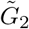 are:

**Figure 7.**
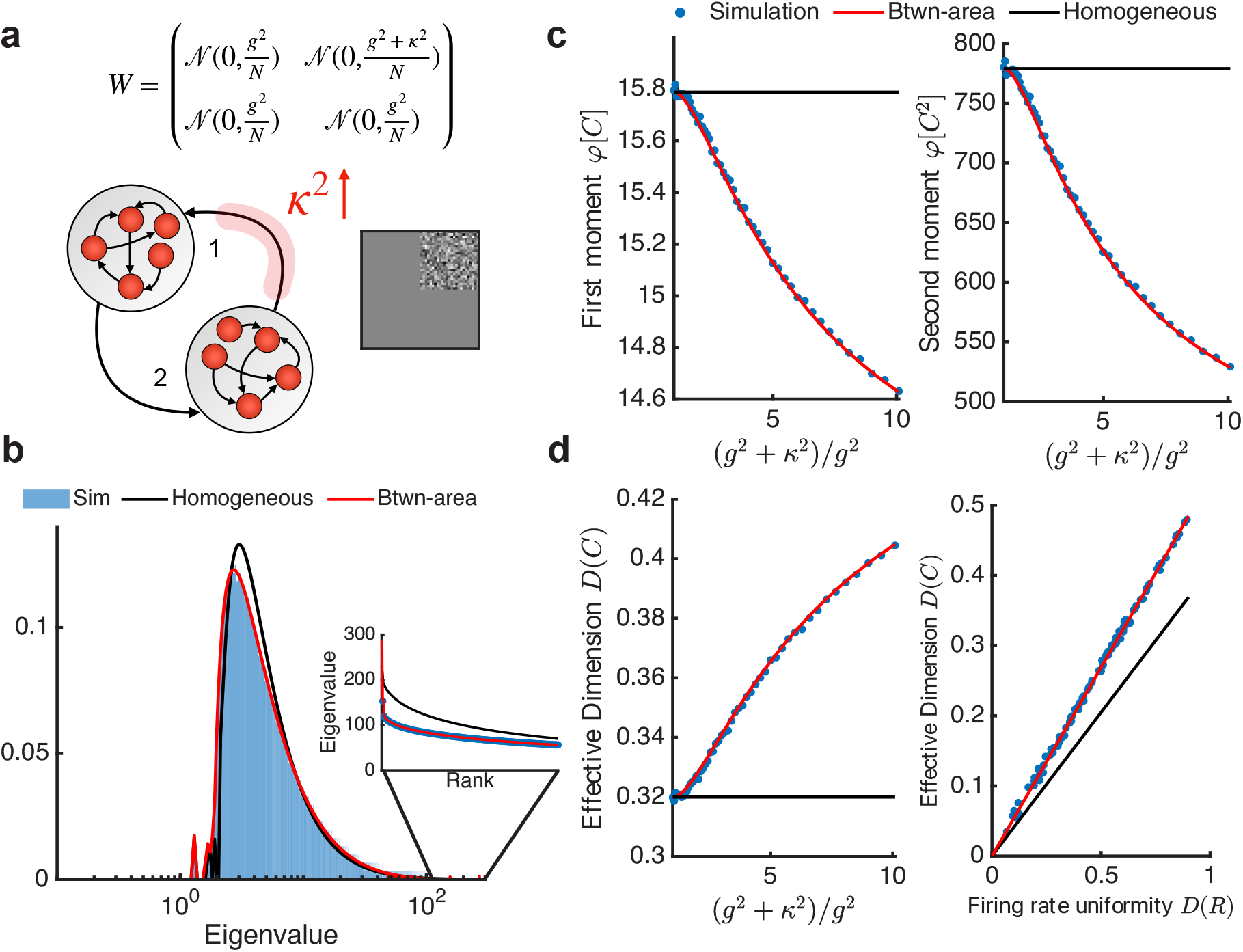
**a)** A recurrent network with two areas and increased variability in connections from area 2 to area 1. The gray square shows the size *κ*^2^ contribution applied to the top right block of the connectivity matrix *W*. *W* is again a 2 *×*2 block matrix, where each block has i.i.d. Gaussian entries. Total variance in *W* is again conserved. **b)** Simulated eigenvalue distribution of covariance matrix *C* against theoretical predictions from Section VII with uniform variance in *W* and Section VIII B with areal-level structure. Theoretical predictions are calculated by solving the self-consistent Cauchy transform equation (Equation 33 vs Equations (47)-(48)) numerically and applying the Stieltjes inversion formula. Solid black line – unstructured connectivity (*κ*^2^ = 0). Solid red line – structured connectivity (*κ*^2^ = 0.11). Inset – largest eigenvalues in the distribution ordered by rank. Blue dots – simulated eigenvalues. Spectral density shows agreement with simulations at both the bulk and the edge. **c)** Theory and simulations of the first and second moments with increasing *κ*^2^. **d)** Left: Theory and simulations for *D*(*C*) across *κ*^2^ match as well. Right: The linear relationship between *D*(*R*) and *D*(*C*) is preserved but the slope increased (*κ*^2^ = 0.11).

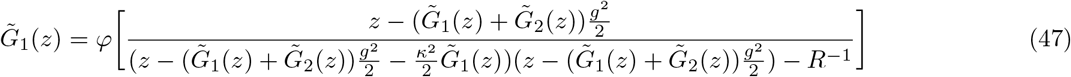

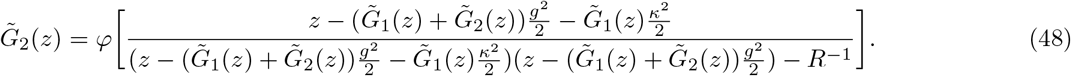

There is excellent agreement between *µ*(*λ*) computed from Equations (47) and (48) and direct numerical estimation from *C* (Figure 7b). Similar to Appendix B, the moments of *C* under this between-area variance structure are:

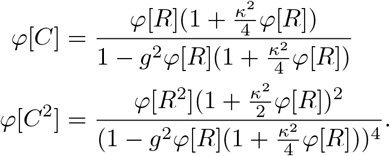

Then, the dimension can be calculated as

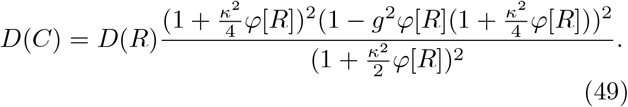

Written in terms of *κ* and reorganized, we have:

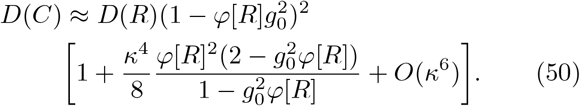

The first and second moments now decrease with increasing between-area variance (theory and simulations shown in Figure 7c), in contrast to the increasing moments from adding variance to within-area connectivity (Figure 6c). Similar to within-area contributions, between-area contributions do not break the linear relationship between *D*(*C*) and *D*(*R*). Instead, the slope increases with added variance (Figure 7d).

Adding variance to the between-area connections from area 2 to area 1, our receiving and sending vectors become:

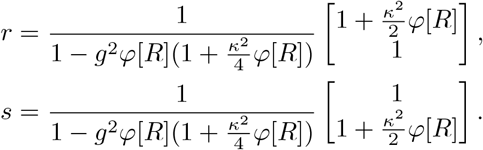

The variance received by area 1 and propagating from area 2 is increased by equal magnitudes, as expected. In contrast to adding within-area variance, adding between area variance separates strong senders from stronger receivers, and the compounding of variance weakens. When the sending and receiving vectors are not as aligned, variance is distributed without concentration, and so decreases by a larger fraction than *φ*[*C*]^2^, ultimately increasing the effective dimension.

### C. Within-area and between-area structures

The effective dimension *D*(*C*) increases with added variance to an off-diagonal block in connectivity (Figure 7), but decreases with added variance to a diagonal block (Figure 6). Taken together, there appear to be opposing forces on the effective dimension depending on how the connectivity variance structure is imposed. To explore this dichotomy we first compare the first moments of *µ* in orders of *κ*^2^. Added within-area structure (after reorganizing Equation (42) up to the fourth order in *κ*) yields:

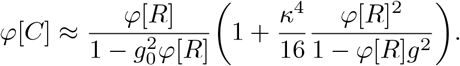

By contrast, increasing between-area connectivity variance has the approximation:

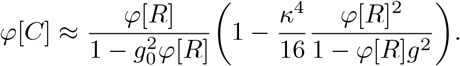

It is clear that the *κ*^4^ terms are identical in magnitude and with opposite signs, so adding or averaging the two may result in an expression that is independent of *κ* (up to *κ*^4^).

In this section we consider added *κ*^2^ along the right block column (Figure 8a). This is equivalent to imposing higher variability in both the recurrent and outward connection weights from area 2. Adding *κ*^2^ along a block row, alternatively, increases variability in recurrent and inward connection strengths. These two cases are mathematically equivalent in the eigenvalue distribution of *C*, differing only in the calculations of 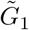 and 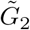.

**Figure 8.**
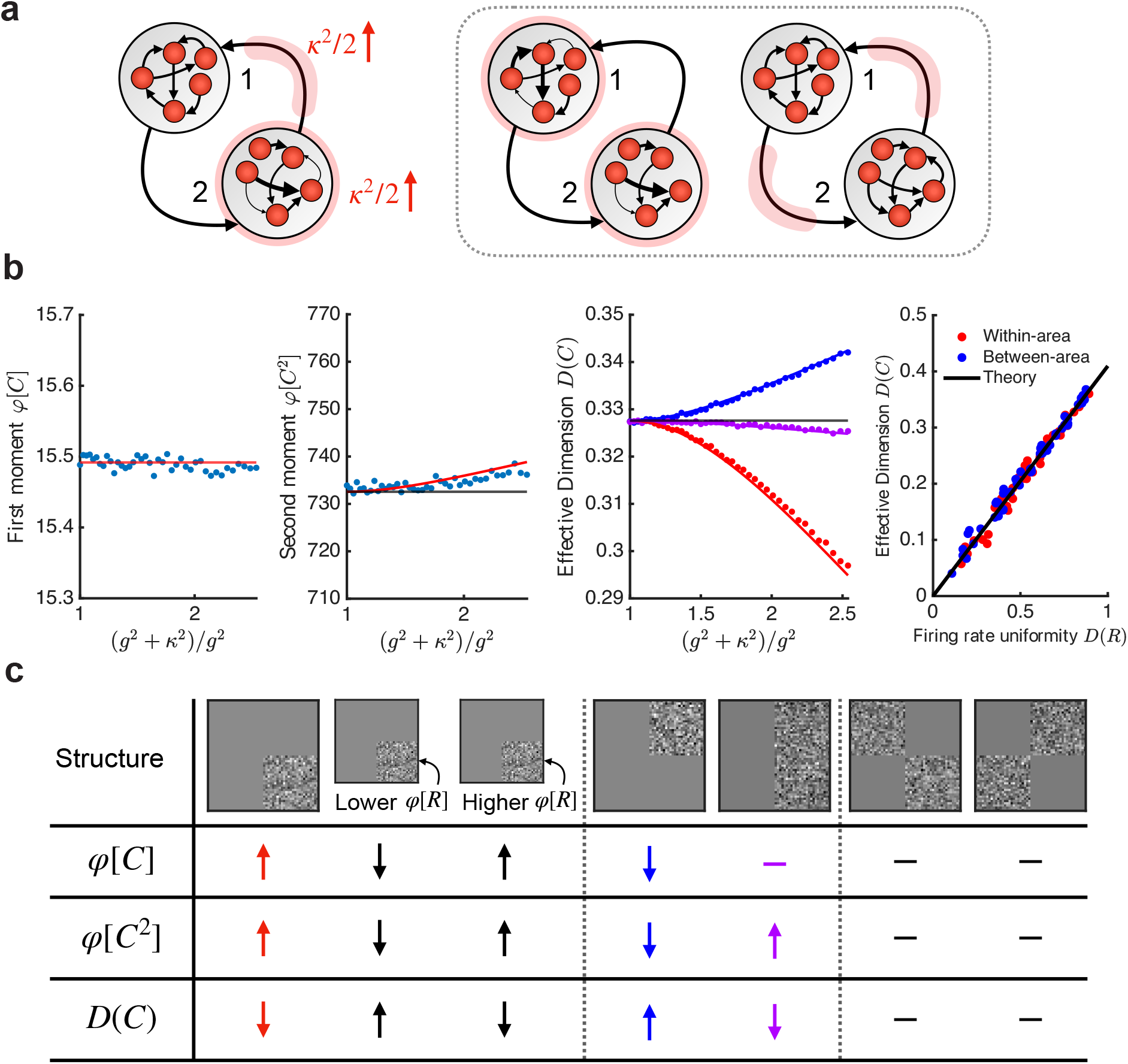
**a)** Schematics showing three ways to add structure with two blocks in *W*. The added variance in each block is of size *κ*^2^*/*2. Left: Increased variance in recurrent and outward connections from area 2, one realization of a simultaneous within- and between-area structure. Dotted gray box encloses two cases where these variance profiles are symmetric across areas. Middle: Increased variance in only recurrent connections within each area (added variance on both diagonal blocks). Right: Increased variance in only between-area connections (added variance on both off-diagonal blocks). **b)** First two panels: Theory and simulations of the first and second moments with increasing *κ*^2^ for simultaneous within- and between-area structure. Solid black line – theory with uniform variance across *W* (*κ*^2^ = 0, Section VII). Solid red line – theory with structure (Section VIII C). Third panel: *D*(*C*) against increasing values of *κ*^2^ across within-, between-, and simultaneous within- and between-area structure. The eigenvalue distribution is balanced with the two types of variance contributions, but the effects from a within-area structure still dominate. Fourth panel: Simulations from symmetric structures with both diagonal blocks (within-area), and both off-diagonal blocks (between-area) against unstructured theory from Section VII. **c)** Summarization table of *φ*[*C*], *φ*[*C*^2^], and *D*(*C*) across all global variance structures. Upward arrows – increase in moments or dimension. Downward arrows – decrease in moments or dimension. Dashes – no change.

We ensure an equivalent amount of added variance *κ*^2^*/*2 to both blocks, so 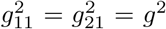 and 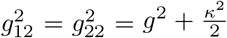. The remaining analysis follows previous calculations (see Appendix B), and we obtain:

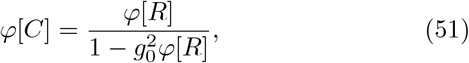

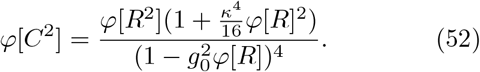

The theoretical and simulated first moments of the eigen-value distribution, *φ*[*C*], do not have a dependence on *κ*, as predicted (Figure 8b, first panel). The second moment, *φ*[*C*^2^], only grows from a small, fourth order term (Figure 8b, second panel).

To observe why the first moment is fixed, we observe that the sending and receiving vectors are:

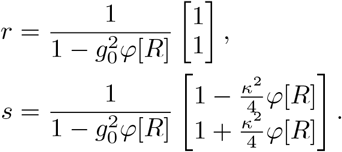

Since *κ*^2^*/*2 was added to the variance in outward connections from area 2, its component in *s* is naturally larger than that of area 1, but their sum is independent of *κ*. In contrast, the areal components in *r* are completely equal. Thus, the balance of the first moment can be interpreted in two ways: equivalent amounts of variability are given to each area (*r*), or the increased propagation of variance in one direction (area 2 to 1) is compensated or diminished by the reverse direction (*s*). Area 1 has lower recurrent variability than area 2 but receives greater variability from area 2, while area 2 has higher recurrent variability but receives less from area 1.

The effective dimension is then:

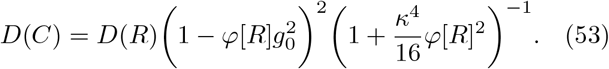

Ultimately, dimensionality decreases despite a balanced first moment. Consequently, despite seeing opposing effects from adding variance to a diagonal vs off-diagonal block, structure imposed on within-area connectivity dominates (Figure 8b, third panel).

It may appear that changes in dimensionality always precipitate from global structures in *W*, but this is not always true. Consider structures that are symmetric across areas, such as increasing within-area variance by size *κ*^2^*/*2 in both areas, or adding between-area variance in both areas (Figure 8a). In both cases, the variance profile *V* is symmetric and is identical in both areas. Both theory and simulations agree that symmetric structures have no influence on the spectral density of *C* (Figure 8b, fourth panel); only asymmetric structures in *W* (Sections VIII A and VIII B) deviate from the unstructured theory in Section VII (Equation 36).

It appears that Equation (36) from Section VII holds when the eigenvalue distribution of *W* matches that of a circular element, with eigenvalues lying on a uniform disk of radius *g*_0_ on ℂ (Figure S1a). To better understand spectral differences in symmetric vs. asymmetric variance profiles, we observe how within- and between-area structures break the circular eigenvalue distribution of *W*. Adding variance to a diagonal block of *W*, we see a larger concentration of eigenvalues within a disk of radius *g* centered around the origin, while the rest of the density becomes more sparse and exceeds the original support of the *W* (Figure S1b). Conversely, adding variance to an off-diagonal block maintains uniformity in the density but reduces the radius of the support (Figure S1c), also diverging from the limiting distribution of the circular law. Together, variance profiles with increased variance in both the off-diagonal and diagonal blocks of *W* offset the radial decrease in density and constrain the support back to the original radius of *g*_0_. However, the bulk is still biased towards an inner radius of *g*. Therefore, not only do we already see the two types of structure counterbalancing each other in the spectral characteristics of the connectivity matrix, we also see the dominance of within-area structure, and that it is more subtle than the non-uniformity observed with within-area structure alone.

Although it is a nice observation that the covariance spectrum is preserved when eigenvalues of *W* follow the circular law, this is not guaranteed due to interaction with *R*^−1/2^. A more rigorous justification is that, with our given symmetric structures, the covariance mapping on the identity is simply a scaled identity: 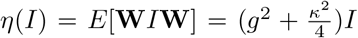(Appendix C). In other words, *η* maps ℂ onto itself. In the unstructured case, this is already evident in Equation (27). With asymmetric variance profiles, an example violation of this relationship is clear in Equation (38). Applying the covariance mapping to the identity matrix with *ℳ*_4_(*C*)−valued element **Ŵ** effectively adds the variance along the block rows and columns of *W*. The first two diagonal elements of *η*(*I*) sum the variance along the two block rows, while the last two diagonal elements sum the variance along the two columns. With summed variance along both the rows and columns being equal in these symmetric structures, variability is uniformly spread across blocks, resembling and thus satisfying the theoretical results for the unstructured network.

### D. Extensions beyond global variance structures in two-areal networks

In Section VII, we saw that increasing total connection variability *g*_0_ decreases *D*(*C*). If we allow the total variance in the network to grow with *κ* or disregard the condition in Equation (37), adding within-area structure strengthens trends seen in the homogeneous case (Figure S2a). Increasing between-area connectivity variance while allowing total variance to grow attenuates the original trends seen in the case where variance is conserved (Figure S2b).

In considering the variance profile *V*, we characterized *φ*[*C*] and *φ*[*C*^2^] in terms of sending and receiving vectors *s* and *r*. An important consequence of this framework is that it enables the generalization of this method to block variances with an arbitrary number of neuronal areas (Appendix D). In general, for *K* areas (same size) we have the compact relation:

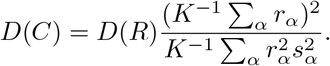

The essential structure is similar to the two area case (Equation (46)).

Finally, in the previous sections we assumed that the firing rates in areas 1 and 2 originate from the same distribution. When neuronal responses from the two areas exhibit different statistical properties, such as different mean firing rates (*φ*[*R*_1_] ≠ *φ*[*R*_2_]), dimensionality may not necessarily decrease with increased variability in within-area connectivity strength. The calculation for *D*(*C*) is nuanced when we need to make a distinction between the firing rates in area 1 (*R*_1_) and the firing rates in area 2 (*R*_2_) (see Appendix E). Holding *D*(*R*) constant, we let *R*_1_ and *R*_2_ be realizations from two log-normal distributions with differing means (Figure S3a). We consider the two cases where area 2 has a lower vs higher mean firing rate than area 1, and again add variance to the connectivity within area 2 (Figure S3b-c). When area 2 has a higher firing rate than area 1 (Figure S3e-f, cyan star), the previous trends are accentuated, and dimensionality decreases more severely with *κ*^2^. However, when area 2 has a lower firing rate (Figure S3d,f, yellow star), added structure instead decreases moments *φ*[*C*] and *φ*[*C*^2^], and increases dimensionality *D*(*C*).

In sum, the effect of a change in *κ* on *φ*[*C*], *φ*[*C*^2^], and *D*(*C*) critically depends on the global structure of both the connectivity variance and the firing rate distributions (Figure 8c). We find that these trends resulting from global structures mirror the effect of local motifs discussed in Dahmen et al. [21], where a higher relative abundance of reciprocal motifs (paralleling localized within-area connections) decreases dimension and more chain motifs (paralleling directed between-area connections) increases dimension.

## IX. SPECTRAL DENSITIES OF INDIVIDUAL AREAS

Current imaging and electrophysiology tools in neuroscience only acquire measurements of a subset of neurons from an underlying population. Ideally, we could study the statistics, such as the covariance, of a subnetwork of neurons and subsequently make inferences about the larger network. In the context of our work, an easy misconception is that the diagonal components of the operator-valued Cauchy transform 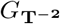, *G*_1_(*z*) and *G*_2_(*z*) (Equation 30), reflect the eigenvalue distributions of individual areas. By a simple consideration of the operator-valued Cauchy transform without structure (*κ* = 0), we can see that *G*_1_ = *G*_2_ = *G*_*C*_. Therefore, *G*_1_ or *G*_2_ cannot possibly provide areal spectral densities, and instead describe covariance globally (Figure 9a). This naturally follows from the understanding that a subnetwork’s spectral density will not be equal to that of the whole network. Instead, *G*_1_(*z*) and *G*_2_(*z*) are eigenvalue projections of 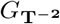 onto each *N* × *N* diagonal block (Appendix F, Figure 9b).

**Figure 9.**
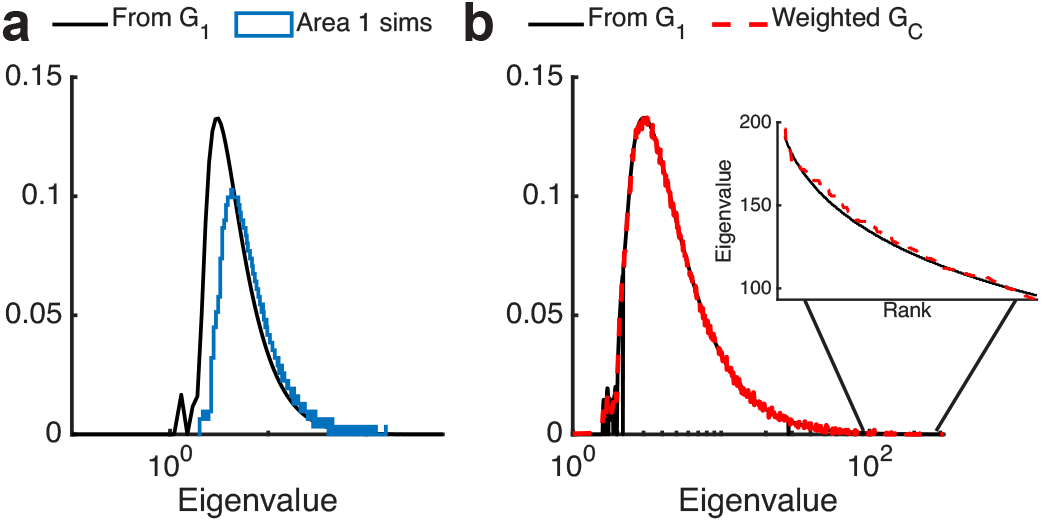
**a)** Eigenvalue distributions derived from a block component of the operator-valued Cauchy transform (*G*_1_) and simulated eigenvalues of area 1. Both simulations and theory have no global variance structure (homogeneous variance, *κ*^2^ = 0), and do not agree. **b)** Areal spectral densities cannot be obtained from *G*_1_, as *G*_1_ is a projection of the operator-valued Cauchy transform onto the top left block of **T**^−2^ (not to be confused with the top left block of *C* or *W*).

Without loss of generality, we will aim to find the Cauchy transform, and thus spectral density, of the covariance in area 1 (top left block). In doing so, we will also take the opportunity to introduce a popular operator-valued free probability tool in finding the limiting measure of polynomials and rationals of large matrices: the linearization trick [24].

### A. The linearization trick

The main idea of the linearization trick is to utilize operator-valued tools to transform nonlinear problems in the algebra into linear problems in the matrices. Polynomials and rationals of non-commutative random variables can be elevated to larger block matrices with a transparent correlation structure, on which we can apply operator-valued free convolutions. Let *P* ∈ ℂ ⟨*X*_1_, …, *X*_*n*_⟩ be the polynomial in the algebra generated by *n* noncommuting variables *X*_*i*_ in *A*, over the field of complex numbers. The distribution of *P* can be deduced from a corresponding *linearization*, written as

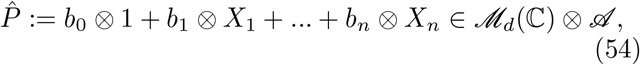

where *b*_0_, …, *b*_*n*_ ∈ *ℳ*_*d*_(ℂ) and *n* is an integer. 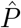 is defined as a matrix given by

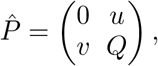

where *Q* ∈ *ℳ*_*d−*1_(ℂ) ⊗ ℂ⟨*X*_1_, …, *X*_*n*_⟩, *u* is a row vector of size *d* − 1, and *v* is a column vector of size *d* − 1, such that 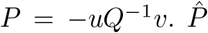 can also be classified as a Kronecker random matrix. For such matrices, self-consistent matrix Dyson equations have been used to approximate the resolvent, from which the spectral density is obtained [99, 100].

As an example, take 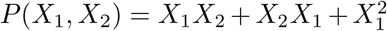 with the linearization

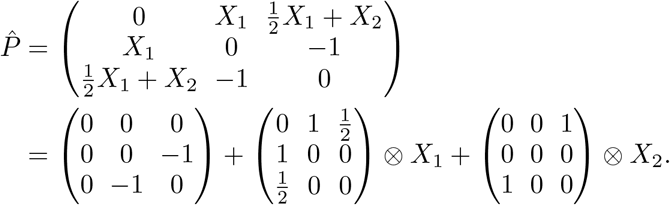

Notice that the entries in *Q* ∈ *ℳ*_2_(ℂ) ⊗ ℂ⟨*X*_1_, *X*_2_⟩, *u*, and *v* are all linear, and 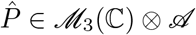 can be broken down into terms of only *X*_1_ or *X*_2_. The linearization trick can be applied regardless of freeness, meaning any polynomial *P* admits a linearization 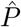. However, we use the freeness of variables for freeness over *ℳ*_*d*_(ℂ), which is necessary for operator-valued free convolutions.

Let us also define

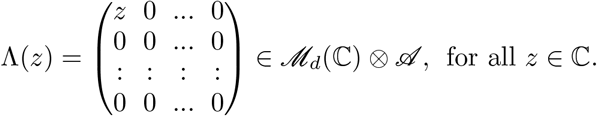

If 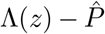 is invertible in *ℳ*_*d*_(ℂ) *A*, then for some *z*, we have

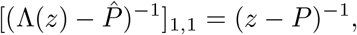

meaning the Cauchy transform *G*_*P*_ can be found by calculating the (1,1) entry of the Cauchy transform 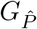:

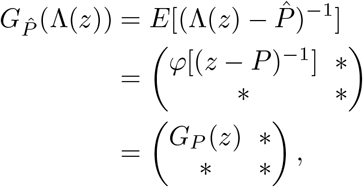

or

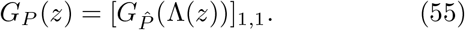

### B. The linearization trick applied to area 1

To apply the linearization trick, we must first obtain a polynomial or rational expression for the top left block of *C* = (*R*^−1/2^ − *W*)^−1^(*R*^−1/2^ − *W*)^*−T*^. Let *R*_11_ and *R*_22_ be the two diagonal blocks of *R*, and *W*_11_, *W*_12_, *W*_21_ and *W*_22_ be the blocks of *W* like before. For brevity, denote 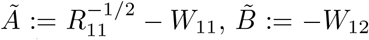,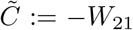, and 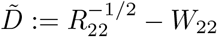.

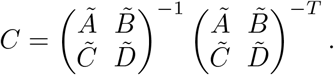

Expanding the expression for *C*:

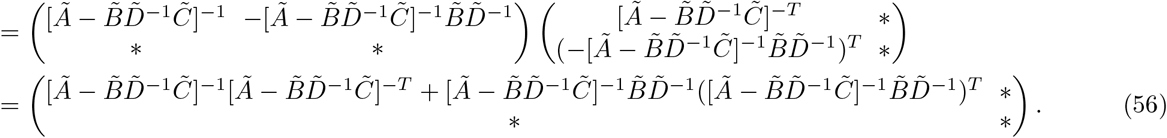

The top left expression of Equation (56) will be our *P* for the covariance of area 1 (the ‘*’ terms are omitted for brevity). In finding the linearization 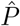, we will break down the long expression into parts, and build the linearization (depicted in Figure 10a). The full linearization 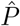 is a 9 × 9 block matrix in terms of *Ã*, 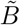, 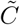, 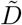, *Ã*^*T*^, 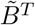, 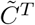, and 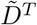, which are all linear expressions of the blocks of *R* or *W*. And so, we can pull all constants and terms from *R* into an *ℳ*_9_(*C*) matrix and all terms from *W* into another *ℳ*_9_(*C*) matrix, call these 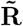 and 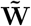 (Appendix G). Since *R* and *W* are free, 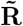 and 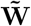 are free with respect to *ℳ*_9_(ℂ), and we can use the subordination property once again in *ℳ*_9_(ℂ). Ensuring that we are in the open set ℍ^+^(*ℳ*_9_(ℂ)), we define Λ_*ε*_(*z*) = diag(*z, iε*, …, *iε*) for *ε >* 0. Then,

**Figure 10.**
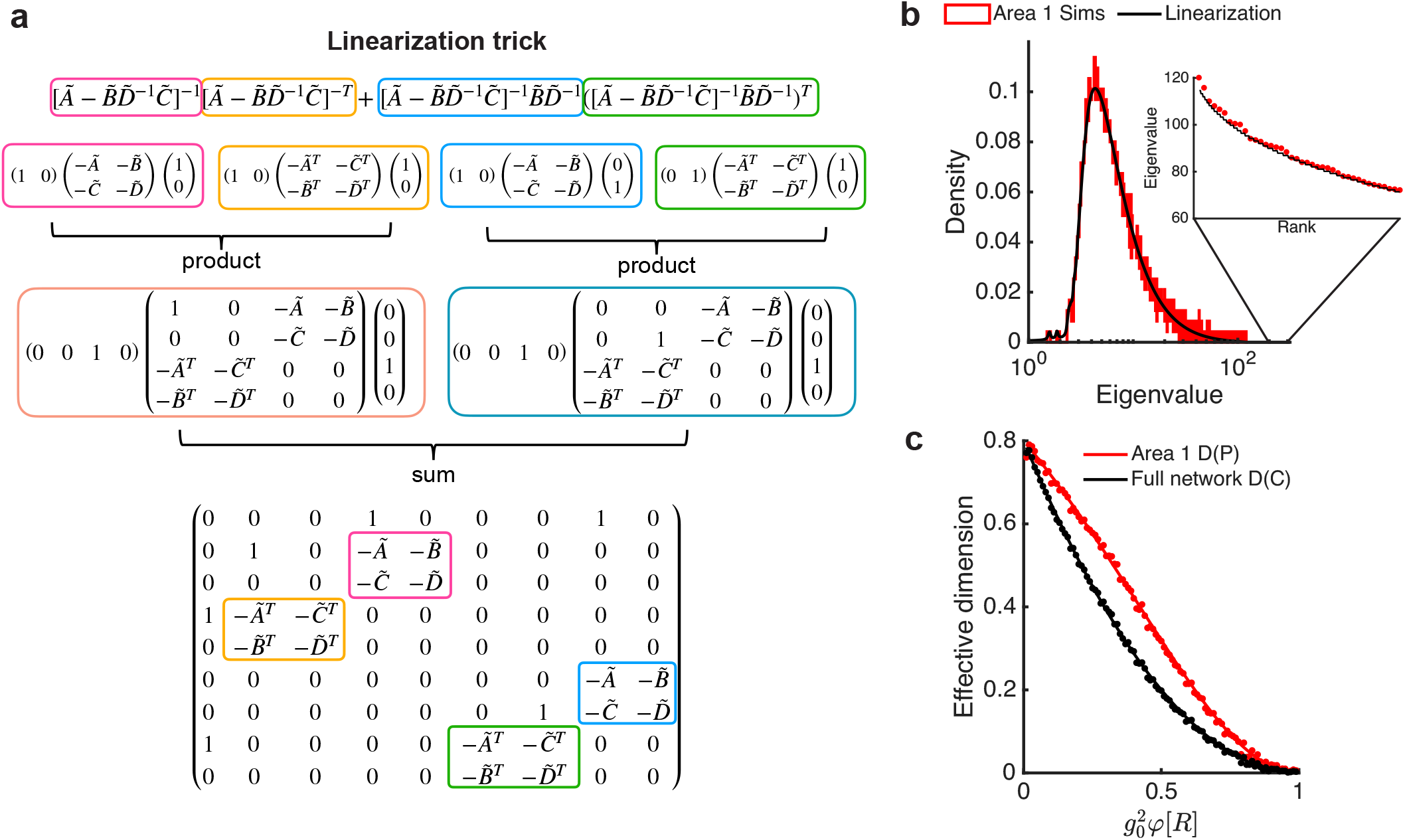
**a)** Visualization of the linearization trick applied to the rational expression for the covariance of area 1 (Expression (56)). The linearizations of smaller components in the rational expression are highlighted in the full linearization through colored blocks. **b)** Numerically solving the Cauchy transform from the full linearization yields an eigenvalue distribution in agreement with simulations. **c)** Simulations and theory for the effective dimension of area 1 (Equation 58) and the full network (Equation 36) match. The term 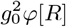 sweeps over the range 0.01-0.99 to ensure stability, but *φ*[*R*] is fixed at 10.

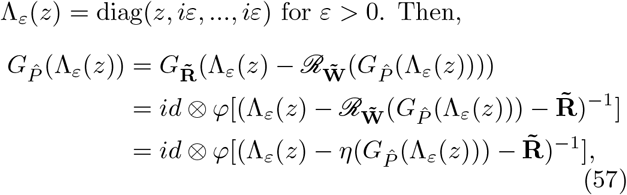

where 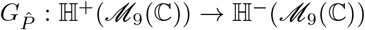. We would then find 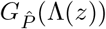 by taking *ε* to zero at the end. Similar to Equation (32), since 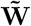 is self-adjoint in an underlying jointly free circular family and their adjoints, 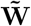 is *ℳ*_9_(ℂ)−valued semicircular. Therefore, we can substitute the *ℛ*-transform 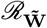 for the covariance map 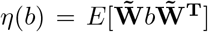, for *b* ∈ *ℳ*_9_(ℂ). By Equation (55), we have

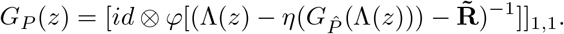

We did not find an explicit solution to *G*_*P*_, but we can numerically solve the system of six equations (see Appendix H) given by entries of the self-consistent Equation (57). The system of equations is excluded from the text due to its length. Taking the (1,1) entry of the 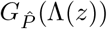 and applying the Stieltjes inversion formula gives the recovered eigenvalue density, which compares well with numerical simulations (Figure 10b).

The normalized, effective dimension for area 1, without any added structure, is

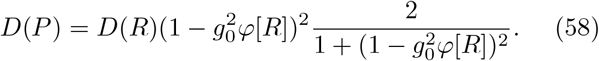

The derivation of the analytical expressions for *φ*[*P*] and *φ*[*P* ^2^], as well as for *D*(*P*) with arbitrary population size *f N* (0 *< f <* 1), can be found in Appendix H. The stability condition requires that 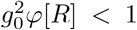, meaning the effective dimension in a single area is always higher (assuming *g*_0_ *>* 0) than that of the multi-areal circuit (Figure 10c).

We did not calculate the analytical expression for *D*(*P*) in a single area with a heterogeneous variance profile. Resorting to numerical simulations, we compare the moments and dimension of individual areas against the full network (Figure S4). Similar to trends of the full covariance matrix described in Section VIII A, adding within-area variance to area 2 leads to an increase in *φ*[*P*] and *φ*[*P* ^2^] of area 2, and an overall decrease in dimension. Alternatively, the moments of area 1 decrease, with its dimension ultimately increasing with *κ* (Figure S4a). The between-area structure from area 2 to 1 shows the converse effect, where moments in area 1 increase and moments in area 2 decrease. However, dimension in individual areas increases along with the whole covariance matrix, though the dimension of area 2 increases more rapidly (Figure S4b). As soon as we split the contribution between blocks (symmetric structures and simultaneous within- and between-area structure), the first moments of each area are also independent of *κ* (Figure S4c-e). With added within- and between-area variance, we once again see dominating within-area structure patterns in that area 1 increases in its dimension while area 2 decreases (Figure S4c). Symmetric profiles along the diagonal blocks show a slight increase in the second moments of individual areas and thus a decrease in their dimensions, but no change in overall dimension (Figure S4d). In contrast, symmetric profiles of off-diagonal blocks show equivalent increases in dimensions in both areas, and again no change in overall dimension (Figure S4e). Even if local statistics in subnetworks change, global statistics do not.

## X. DISCUSSION

In this work, we studied how heterogeneity in the connectivity of recurrent networks shapes the effective dimension of activity covariance. We considered both local and global levels of heterogeneity in connection weights between neuron pairs, as well as cellular heterogeneity as measured by firing rates. The former has been discussed extensively in our previous work in local heterogeneity [23]; its derivation and an extension to global heterogeneity are presented here using various tools of operator-valued free probability. We found that the dimension of covariability decreases with added within-area connectivity variance, but increases when this added variance is in connections between areas. We considered when these structures are able to balance eigenvalue statistics, distinguishing between symmetric vs asymmetric variance profiles across areas. We also introduced tools to handle functions of random matrices, obtaining eigenvalue distributions of individual areas within the network.

### A. The relation between *D*(*R*) and *D*(*C*) in cortical data

For unstructured networks, we derived [23] a linear relationship between *D*(*R*) and *D*(*C*) (Equation 36). Analysis of simultaneously recorded neuronal populations in sensory, cognitive, and motor regions in mice, monkeys, and humans tested this prediction. This success in predicting an observed connection between the distribution of mean responses and the effective dimension of population variability was the central result of our past work.

While this agreement between theory and experiment is satisfying, the assumption of unstructured random wiring between neurons ignores much of the known circuitry of cortical networks, where neurons are often pooled into regional or functional groups [29–33]. A natural question to ask is: when these biological details are considered will the linear relationship between *D*(*R*) and *D*(*C*) vanish? If the linear relationship was dependent on a lack of structure (i.e. freeness of *W* and *R*) then its validation in data might suggest that any structured aspects of recurrent coupling in real cortical networks were negligible for simple statistics such as *D*(*C*). This conclusion would be unfortunate, as it suggests that structure in cortical wiring may be less consequential (yet see [21]). Fortunately, in this study we established that the linear *D*(*R*) and *D*(*C*) relationship extends to networks with global structure in *W*, and we further understand how global heterogeneities affect the linear relationship between *D*(*R*) and *D*(*C*). In particular, an attractive application of block structure in *W* could be to consider excitatory and inhibitory synaptic wiring in the network. In total, the validation of our past theory in data does not require an unrealistically simplistic model of unstructured cortical wiring.

Nonetheless, in our previous study [23] we showed how the linear relationship between *D*(*R*) and *D*(*C*) breaks down when populations are composed from simultaneous recordings of neurons spanning two brain regions. While differences in the between and within brain-areas connectivity are well established, the above discussion shows that the lack of a linear relation between *D*(*R*) and *D*(*C*) cannot be explained by simply a global block structure in *W*. Other alternatives, such as a low rank connectivity structure between brain areas or sources of external correlation (i.e. ⟨*ζζ*^*T*^⟩ ≠ *I*), should be next considered to explain the breakdown of the linear relationship between *D*(*R*) and *D*(*C*).

### B. Connections to other work

In contrast to our work, Clark et al. [27] found that the dimension of activity increases with connectivity variance when networks have unstructured (i.e. local) connectivity. However, their network has strong recurrent connections, so dynamics are chaotic and variability is internally generated, while our framework inherits and alters variability from external inputs (i.e., from ⟨*ζζ*^*T*^⟩). They developed a two-site cavity dynamic mean field theory to calculate (cross-)covariances and incorporated a symmetry structure through coupled cavity units, which requires the structure in *W* to be fully reflected in the local coupling of the units. While the authors did not analyze other network-wide structures, they discuss how an expansion of their framework can handle nonlocal structures involving within-area and between-area coupled statistics, as shown here, but likely not other global constraints like orthogonality. If we assume orthogonal matrices are pulled from a uniform distribution over the orthogonal group, i.e. Haar-distributed orthogonal matrices, applications of free probability tools directly follow, as independent Haar-distributed orthogonal matrices are asymptotically free from other random or deterministic matrices [24, 101, 102]. We do not complete this analysis, but we note the favorable property that the normalized traces of powers of Haar orthogonal random matrices go to zero, and that the matrices converge to Haar unitaries in non-commutative probability spaces [24, 103]. Therefore, our framework could theoretically replace *W* with an independent Haar orthogonal matrix, and those calculations in the large *N* limit will only lead to changes in the *ℛ*-transform and covariance calculations.

Clark et al. [27, 28] show examples where additional structure beyond i.i.d. connectivity decreases dimension. The study [27] required sufficient symmetry structure, while the latter [28] added constraints on the rank of the connectivity matrix. Our work elucidates cases where global structure in connectivity can actually increase dimension: *D*(*C*) increases with added connectivity variance between areas of the network (Figure 7) and even when variance is added within an area, as long as it has a comparatively lower mean firing rate (Figure S3).

Dahmen et al. [21] found that dimensionality increased with additional connectivity structure, but specifically when that structure took the form of local, second-order motifs (convergent, divergent, chain) that decreased network recurrency, defined as the spectral radius of connectivity eigenvalues (our 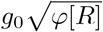). Recanatesi et al. [26] similarly calculated the participation ratio of the covariance in terms of second-order motifs, but found the opposite – that increasing most motifs decreases dimensionality. Our multi-areal network can also incorporate local connectivity motifs if we construct *W* as a sum of real Ginibre ensembles and rank-one random matrices, as detailed in the appendix of [21], with additional block structure.

### C. Selecting intermediate subalgebras

As we have seen through the hermitization of *W* in Sections VIII A-VIII C and the linearization trick, raising random variables and functions of those variables to the operator-valued setting simplifies many problems. Tools used in the operator-valued setting are direct extensions of tools in the free probability setting (e.g. conditional expectation, Cauchy transform, *ℛ*-transform, subordination), rendering application rather simple. Semicircularity and circularity of operator-valued elements also serve as clean extensions from free probability. The former has utility in reducing the *ℛ*-transform and restricting the covariance map, and both have non-trivial, but straight-forward relationships to scalar-valued semicircularity and circularity. As such, the crucial aspect of operator-valued distributions and techniques is that everything is with respect to an intermediate subalgebra.

As discussed throughout this work, if operator-valued elements are in algebra *A*, the conditional expectation is defined according to subalgebra *ℬ* ⊂ *A*, where *E* : *A* → *ℬ*. Selecting *ℬ* appropriately becomes critical, as freeness is consequently also defined with respect to *ℬ* For example, two block matrices in *ℳ*_2_(*C*) that are free with respect to *ℳ*_2_(ℂ) may not be free in the scalar-valued non-commutative probability space (*ℳ*_2_(*C*), *tr* ⊗ *φ*). Furthermore, operator-valued elements that are *ℬ*−valued semicircular may not be scalar-valued semicircular. Generally, semicircularity depends on the level of the intermediate subalgebra, with preservation of semicircularity only in special cases. Sections VIII A-VIII C depict one example of such cases where semicircularity is passed from *ℬ* to *D*, the subalgebra of diagonal block matrices. We then note that the careful selection of the intermediate subalgebra and the necessarily discrete nature of matrix blocks hinder the ability to build connectivity structures with gradients (Figure 2 in [23]).

### D. A nod to standard physics-based RMT methods

Taking the semicircle law as an example, there are numerous approaches to recovering a spectral density: Coulomb gas techniques [63, 64, 104], analytical approaches that utilize the resolvent [61], combinatorial proofs or the moments method [61], large deviation theory [24, 63, 104], the replica trick [19, 68, 105, 106], and the cavity method [68, 107]. The last two methods, replica and cavity, are ubiquitous in physics texts. The replica trick is suitable when the problem of interest is high dimensional, has disorder whose averages are analytically tractable, and involves finding a self-averaging statistic. The self-averaging statistic in conventional random matrix theory problems is typically the spectral density, and averaging over disorder involves averaging over matrix ensembles. Recovering the density in these scenarios either involves the logarithm of a partition function, averaged over the relevant matrix ensemble, or the logarithm of the characteristic polynomial (Figure 3b). Hu and Sompolinsky [19] showed, with the replica trick, the same results from Section VII (Equation 36) with homogeneous gain (so did [54] and [41]). However, because the replica trick is based on heuristic arguments rather than rigorous mathematics, its results typically must be validated post hoc.

Alternatively, the cavity method is generally more rigorous and can give exact asymptotic solutions for models of disordered systems. Given a large network, it assumes that if one node is added or removed in the network, the remaining nodes become approximately conditionally independent in the large limit *N*. Observing the effect of adding a new site, such as the resolvent under a new row and column [69] or the Gibbs-Boltzmann distribution from the Edwards-Jones formula (Figure 3b) after removal of a node [68], one can derive self-consistent equations to calculate the Cauchy transform. The cavity method contains more concrete interpretability, has direct algorithmic connection, and provides local information in the network, but is also more difficult to manipulate. Although it is likely that the cavity method can produce results similar to those in this work, we do not resort to those methods.

Cortical neurons are rather sparsely interconnected, with a single neuron receiving direct synaptic input from less than 3% of its neighbors [108]. Sparsity, such as with sparse random graphs, tests the universality of spectral densities because the distributions of individual entries change with matrix size [109]. Sparse non-Hermitian matrices with zero mean and unit variance obey the circular law [72], while sparse normalized adjacency matrices of an Erdős-Rényi graph obey the semicircular law [110, 111] (accounting for deterministic shifts in fluctuations at the spectral edge [112]). However, both laws only hold when the number of nonzero entries per column/row tends to infinity (local semicircular law is only obtained under a certain rate: ≥ (log *N*)^*C*^ for some constant *C* and matrix of size *N* [110, 111]; the circular law has no such constraint [74]). If these conditions are satisfied, the nice properties of semicircular and circular elements in free probability calculations likely follow. If not, replica and cavity methods provide more flexibility in determining limiting spectral densities, such as a generalized circular law [25, 113] or a set of numerically solved equations that can recover the semicircle law in the dense limit [68, 113– 115]. This work only presents dense ensembles that are necessarily full rank.

## DATA AND CODE AVAILABILITY STATEMENT

All code used for this work can be found at https://github.com/julietj135/dimensionality-global-heterogeneity.git.

## ACKNOWLEDGMENTS

B.D. is supported by the NIH grants R01EY037119, R01NS133598, T90DA059109; NSF grant 2605555, and the Simons Foundation Collaboration on the Global Brain. J.J. is supported by the NSF Graduate Research Fellowship Program.

This research benefited from Physics Frontier Center for Living Systems funded by the National Science Foundation (PHY-2317138). Financial support was provided via the National Institute from Mathematics and Theory in Biology (Simons Foundation award MP-TMPS-00005320 and NSF award DMS-2235451).

This work was supported in part by a grant of access to OpenAI models through the ChatGPT for Academic Researchers program. Some calculations from Appendix B and H benefited from discussions with Astra.

## Appendix A: Numerical solvers and simulations

In all figures, firing rates *R* are drawn from a log-normal distribution ℒ*N*(*µ, σ*^2^), with *µ* and *σ*^2^ chosen for a given *D*(*R*) and *φ*[*R*]. In the simulations of Figure 5b (black dots), for example, we varied *D*(*R*) by changing *µ* and *σ* while *φ*[*R*] and *g*_0_ were fixed. In contrast, the theoretical plots had *g*_0_ fixed and *φ*[*R*] increasing. We add here that our result in Section VII, Equation 36, does not depend on the exact distribution of firing rates *R* (see Fig. S2 of [23]). For this reason, our work does not test different firing rates distributions.

In Figures 5a, 6b, 7b, 9a-b and 10b, each system of equations for the Cauchy transform was solved numerically using MATLAB’s fsolve function with fixed *R*. For a system of *k* equations, the vector [−0.1*i*, …, − 0.1*i*] ∈ [ℂ^*−*^]^*k*^ was used as an initial guess. If the resulting density shows numerical failures, the magnitude of initial guesses in ℂ^*−*^ was increased. Application of the Stieltjes inversion formula occasionally resulted in negative density values that indicated ether convergence to a non-physical root near the spectral edge or a branch-cut artifact within the bulk.

For the bulk, we noticed that the magnitude of these values followed continuously from neighboring points, so we took the magnitude to correct for the artifact. For the edge, it is possible that the negative density root is genuine but is nevertheless close in magnitude to the true physical root, or other numerical instabilities, so again we simply took the density magnitude at this point. After these corrections, we observed that the resulting density maintains its smoothness and also agrees with simulations.

## Appendix B: Derivation of the moments and effective dimension of covariance in networks with globally structured connectivity

When adding within-area variance *κ*^2^ to area 2 of *W*, the Cauchy transforms 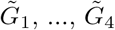 have the relationships:

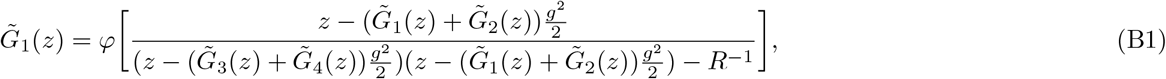

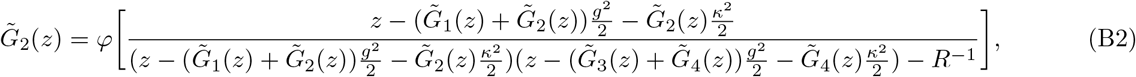

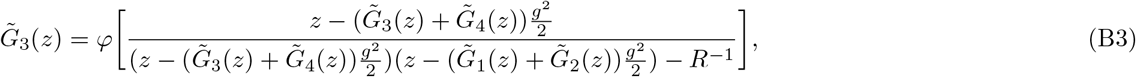

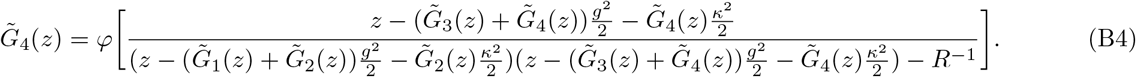

These equations were reduced to Equations (39)-(40) in the main text for brevity and comparison against the homogeneous case, but we include all four here in the Appendix to set up a more general calculation. Let us define

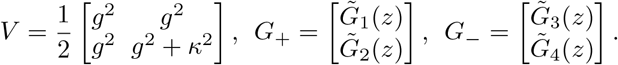

In defining the connectivity variance profile *V*, generalizing to any connectivity structure only involves changing the entries of this profile. Then, the system of four equations becomes:

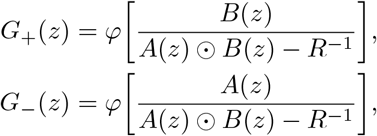

where

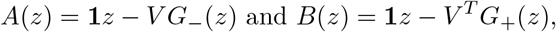

and ⊙ denotes a Hadamard product.

As explained in Section VIII A of the main text, we are trying to find the *z* and *z*^3^ terms of

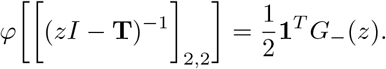

To solve for terms of *G*_*−*_(*z*), we expand the argument of *φ*[·]:

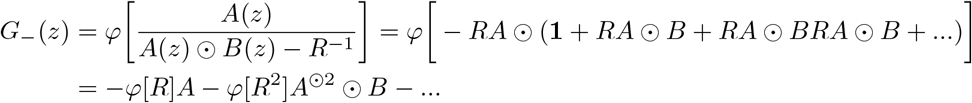

and do the same for *G*_+_(*z*). To find *φ*[*C*], we seek only terms of *G*_*−*_(*z*) that are linear in *z*. Then,

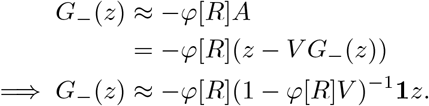

Let *r* := (1 − *φ*[*R*]*V*)^−1^**1** for the sake of brevity, so *G*_*−*_(*z*) = −*φ*[*R*]*rz* + *O*(*z*^3^). This means the linear term of 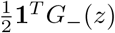 is

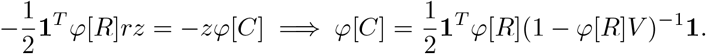

Rewriting the expression for *φ*[*C*] in terms of the two components of *r*, we have:

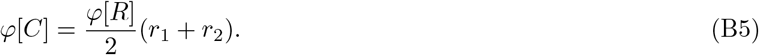

Additionally,

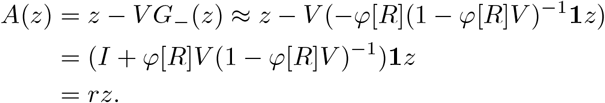

And so, if we let *s* = (*I* − *φ*[*R*]*V* ^*T*^)^−1^**1**, then we also have

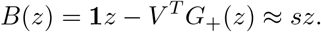

For the second moment *φ*[*C*^2^], we would need to find the *z*^3^ term. Let us introduce a new 2 × 1 vector *h* such that

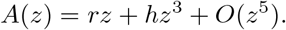

We now solve for *h* by extending the expression for *G*_*−*_(*z*) to the next degree:

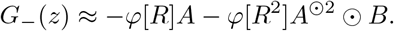

Plugging this expression for *G*_*−*_(*z*) back into the expression for *A*(*z*), we have

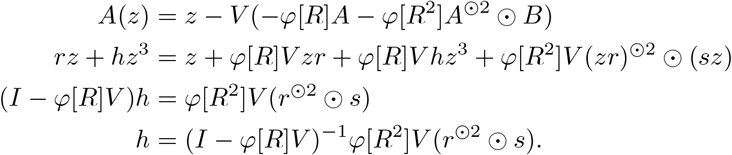

So now we know the *z*^3^ term of *A*(*z*), and can find the *z*^3^ term of *G*_*−*_(*z*):

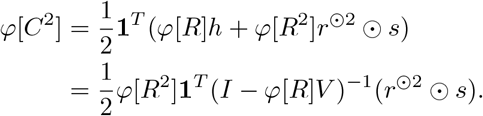

The expression for *φ*[*C*^2^] rewritten in terms of components of *r* and *s* is:

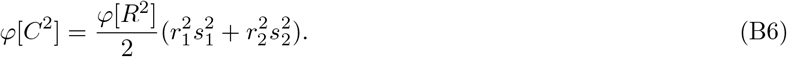

Together, we get:

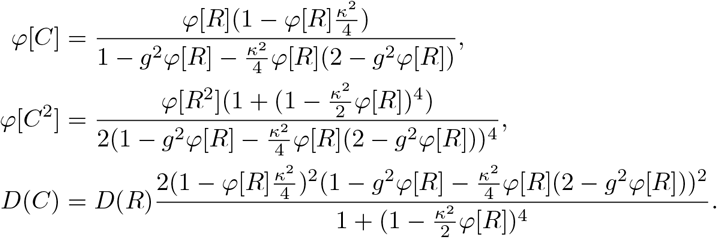

## Appendix C: Symmetric profiles spread variance evenly

Here we show the quick calculation of the covariance mapping for symmetric variance profiles. When both diagonal blocks have added variance, we have

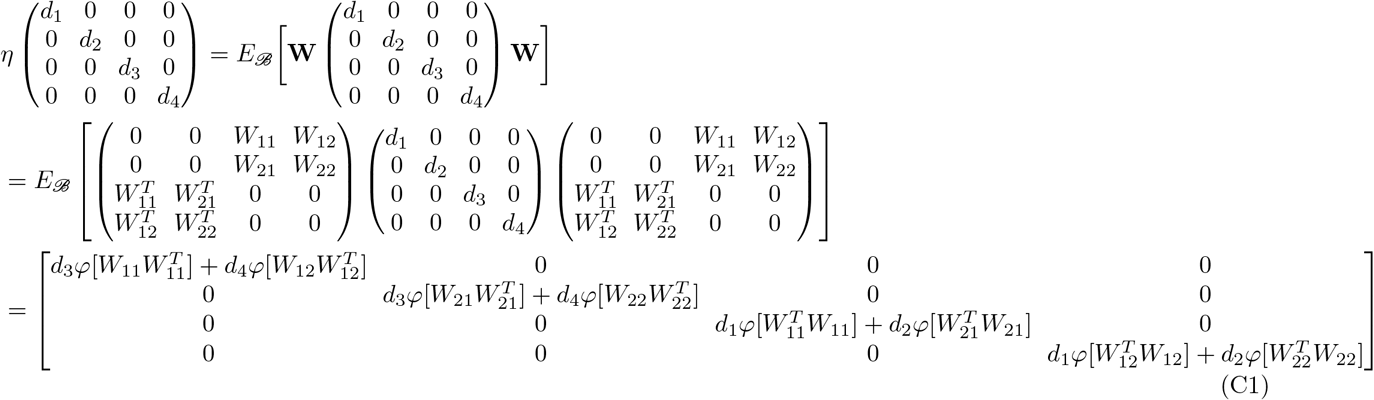

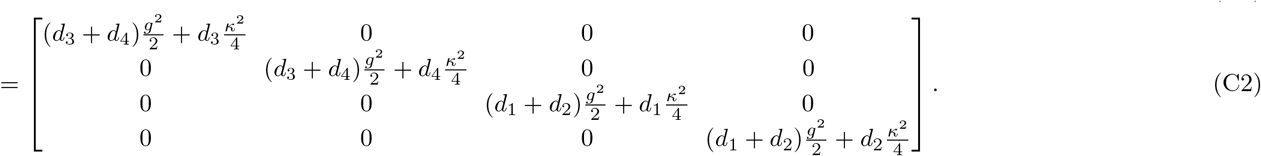

When both off-diagonal blocks have added variance, this is

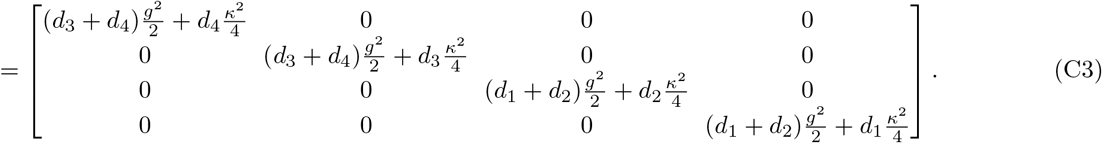

In either Expression (C2) or (C3), 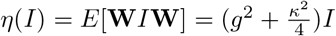. Expression (C1) illustrates how each entry along the diagonal is either the sum of variances along a block column or row if the input to *η* is the identity matrix.

## Appendix D: Generalization of *φ*[*C*],*φ*[*C*^2^], and *D*(*C*) to *K* areas

In Appendix B, *V* is a 2 ×2 matrix and *G*_+_ and *G*_*−*_ are 2 ×1 vectors for a network with two areas. Taking into account the dimension of these variables, this framework can easily be extended to an arbitrary number of *K* areas of equal size. The only distinction in the calculations comes in the last few steps of finding *φ*[*C*] and *φ*[*C*^2^]. With *K* areas, we define *V* ∈ *M*_*K*_(ℝ^+^) again as the variance profile of the network with entries 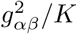, and *G*_+_ and *G*_*−*_ as *K* × 1 vectors. Then, we have

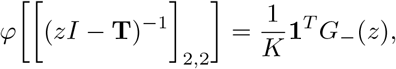

so

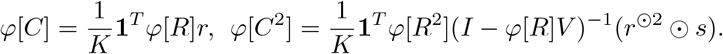

Notice how the expressions have virtually no changes; the extension simply entails averaging contributions over a greater number of areas. Putting the above together yields:

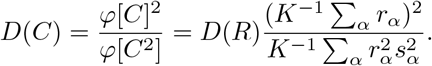

## Appendix E: Calculations for networks with different firing rates

Let area 1 have firing rates *R*_1_, area 2 have firing rates of *R*_2_, and the *κ*^2^ contribution be within area 2. **Ŵ** does not change from Section VIII A, but now

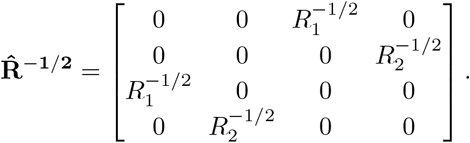

Even when making distinctions between *R*_1_ and *R*_2_, the equations for 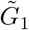 and 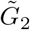 of *G*_**T**_ look almost identical to Equations (39)-(40):

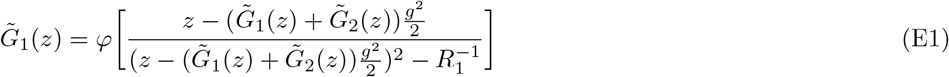

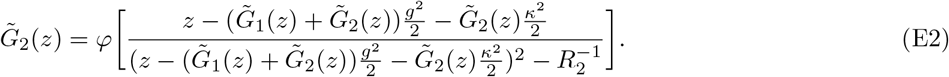

Following the procedure detailed in Appendix B, the first and second moments are

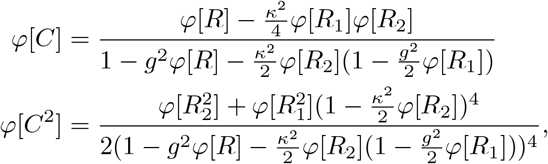

with 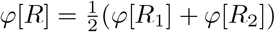. The participation ratio is

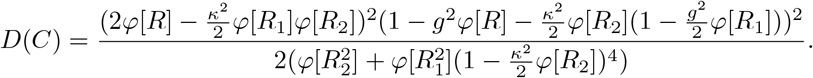

Theory and simulation agreement for moments and *D*(*C*) is shown in Figure S3.

## Appendix F: Derivation of *G*_1_ and *G*_2_ as projections

Recall that *G*_1_ and *G*_2_ are the diagonal block components of 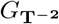, the operator-valued Cauchy transform of **T**^−2^. By the definition of an operator-valued Cauchy transform and the definition of a conditional expectation,

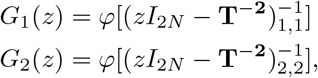

where each argument of *φ* is a diagonal block of the resolvent of **T^−2^**. Using spectral decomposition, we can write the resolvent as

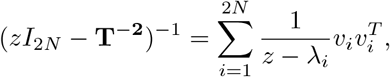

where *λ*_*i*_ is the *i*th eigenvalue of **T**^−2^ and *v*_*i*_ is the corresponding eigenvector. Then,

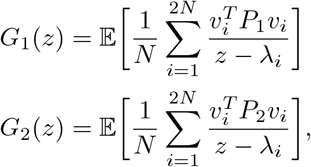

with

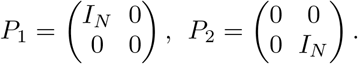

## Appendix G: Linearization of area 1

The 9 × 9 matrix in terms of *A, B, C*, and *D* is shown in Figure 10a. Here we explicitly write out the matrices 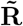 and 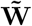, as they are too large to fit in the main text:

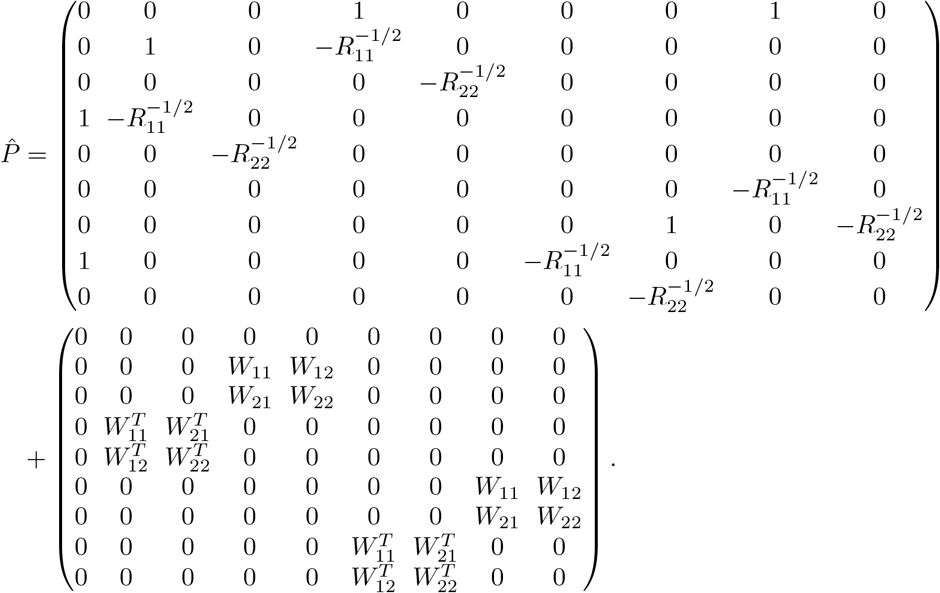

## Appendix H: Deriving the effective dimension of area 1

Recall from Equation (9) that the Cauchy transform is a type of moment generating function. The first and second moments of *P* are then the *z*^−2^ and *z*^−3^ terms of *G*_*P*_ (*z*) or 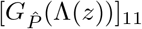. We will solve the self-consistent equation given by the subordination property applied to 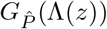 (Equation 57) to the necessary orders, from which we can obtain 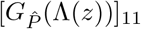. 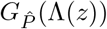 is a 9 × 9 block matrix, and we will let *g* = *g*_0_ for this unstructured calculation. Writing the large matrix as {*G*_*ij*_} for convenience and calculating 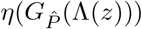 gives:

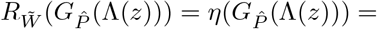

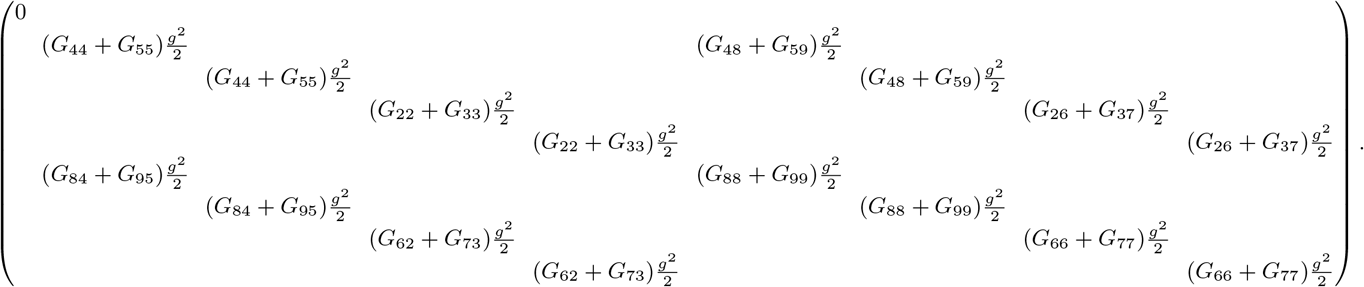

Redundant terms and symmetry reduce this large system of equations to just six variables. Let those six variables be written as *G*_2233_ = *G*_22_ + *G*_33_, *G*_4455_ = *G*_44_ + *G*_55_, *G*_6677_ = *G*_66_ + *G*_77_, *G*_8899_ = *G*_88_ + *G*_99_, *G*_2637_ = *G*_26_ + *G*_37_ = *G*_62_ + *G*_73_, *G*_4859_ = *G*_48_ + *G*_59_ = *G*_84_ + *G*_95_, and define the vector

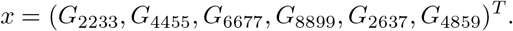

It is important to note that these variables are not Cauchy transforms on their own. As *z* goes to infinity, Cauchy transforms should be on the order of *z*^−1^. While this is true for 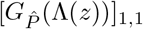, it can be checked by calculating 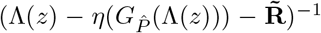 v that the expressions for some variables have leading order *O*(1) rather than *O*(*z*^−1^). Therefore, we will write *x*(*z*) = *x*_0_ + *z*^−1^*x*_1_ + *O*(*z*^−2^).

Defining *u* = (0, 0, 1, 0, 0, 0, 1, 0)^*T*^, write

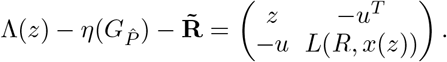

Denoting [·]_8_ as the bottom right 8 × 8 block of a larger 9 × 9 block matrix, we have

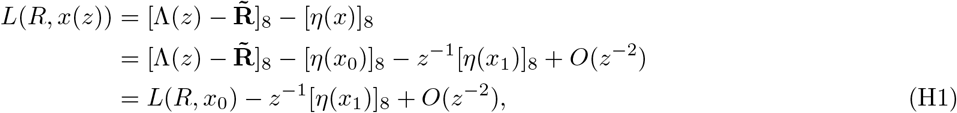

because *η* is linear. Using the Schur complement, we can separate the (1, 1) entry of 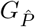 from the rest of the matrix:

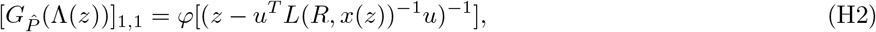

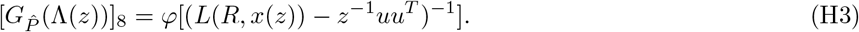

Expanding the expression in *φ* for 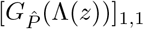 gives

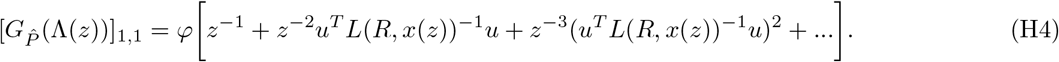

Therefore, we must solve for *x*_0_ and *x*_1_ to properly obtain the *z*^−2^ and *z*^−3^ terms of this expansion. To calculate *x*_0_, we utilize the definition of the Cauchy transform rather than the subordination property:

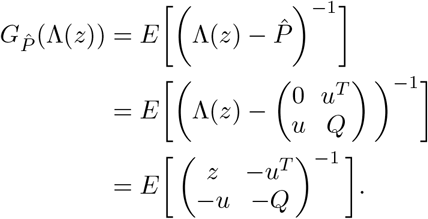

All entries of *x* are in 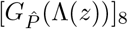, so we only need the bottom right block of this inverse:

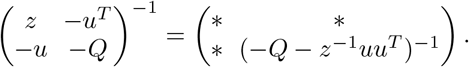

Expanding the bottom right expression, we have (−*Q − z*^−1^*uu*^*T*^)^−1^ = −*Q*^−1^ + *O*(*z*^−1^). Taking *z* to infinity to find constant terms *x*_0_, we only need to observe entries in *Q*^−1^. From Figure 10, it is clear that

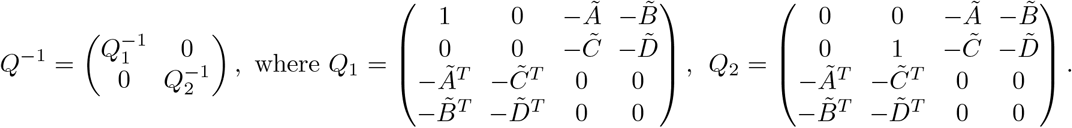

Observations reveal that the constant terms of *G*_22_ and *G*_33_ are in the upper left block of 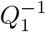, which are zero. Thus, the first term of *x*_0_ is zero. Similar conclusions can be drawn for *G*_6677_, *G*_2637_, and *G*_4859_, leaving two nonzero entries of *x*_0_. To find the constant terms of *G*_4455_ and *G*_8899_, we can re-calculate *id* ⊗ 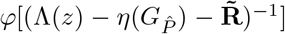 with only the constant terms of 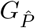, or only *x*_0_. Since most of these terms are zero, the self-consistent Equation (57) ultimately gives

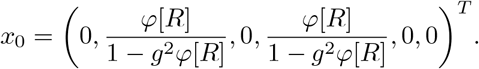

To find *x*_1_, we expand the rational expression for 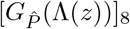 (Equation H3) while substituting the expansion of

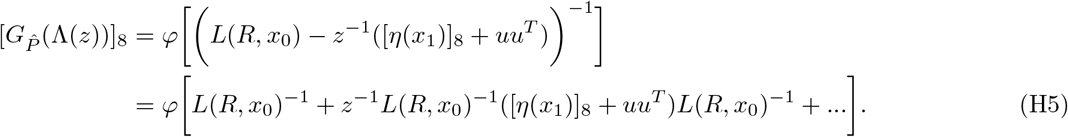

We can easily find *L*(*R, x*_0_)^−1^ because *x*_0_ is known. Given that the *x*_1_ term in Equation (H5) is linear, solving for *x*_1_ is only a matter of matching terms on both sides of the equation. Doing so gives

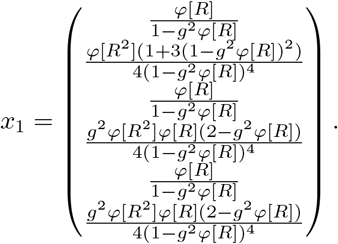

Going back to Equation (H4), we can now expand *L*(*R, x*(*z*))^−1^ as

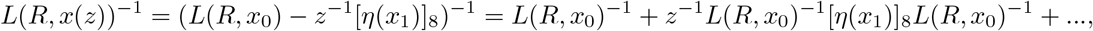

and plug this expansion into 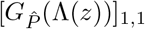:

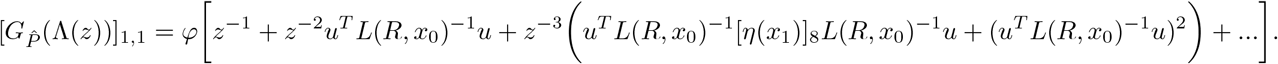

Matching terms with the series expansion of 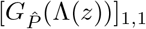, we have

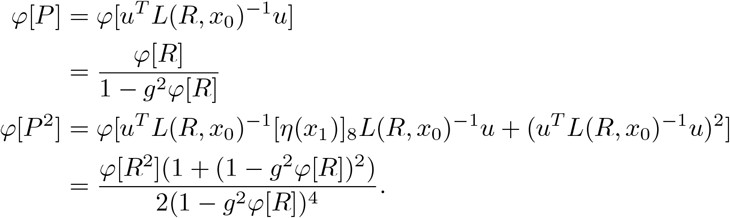

Consequently, the effective dimension of the covariance in area 1 is

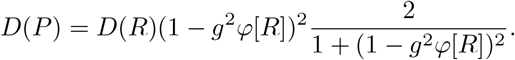

For arbitrary area sizes of fraction *f*, 0 *< f <* 1, rational expression *P* for the covariance in area 1 now has size *f N* × *f N*, while that of area 2 has size (1 − *f*)*N*. The calculations remain the same, except the covariance mapping 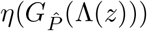 now has weighted sums instead. The six variables of interest must also be updated to include these weighted sums:

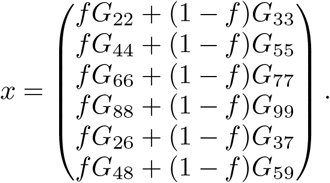

Writing *x*(*z*) = *x*_0_ + *z*^−1^*x*_1_ + *O*(*z*^−2^), we get

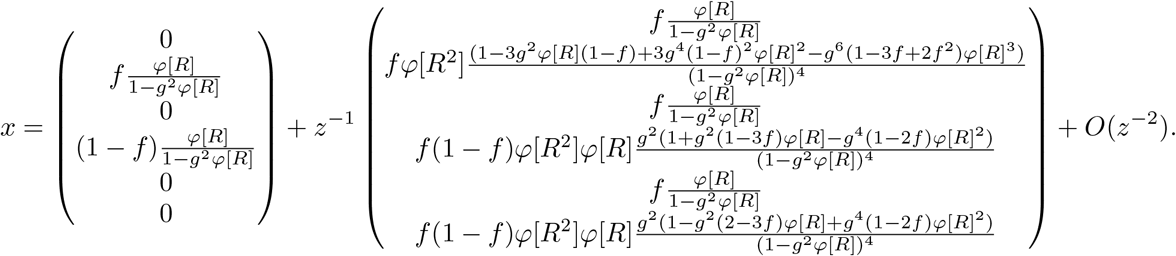

Therefore,

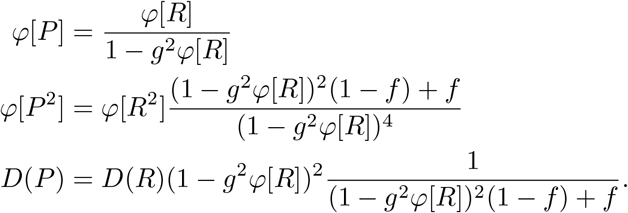

When *f* = 0.5, this is equal to Equation (58). Increasing the fraction *f* of neurons sampled monotonically decreases the effective dimension. In other words, without full access to the network, the observed normalized dimension is always an overestimation. Hu and Sompolinsky derived identical results for spatially sampled covariance matrices with homogeneous gain (*L* = *I*) [19].

---

**Figure S1.**
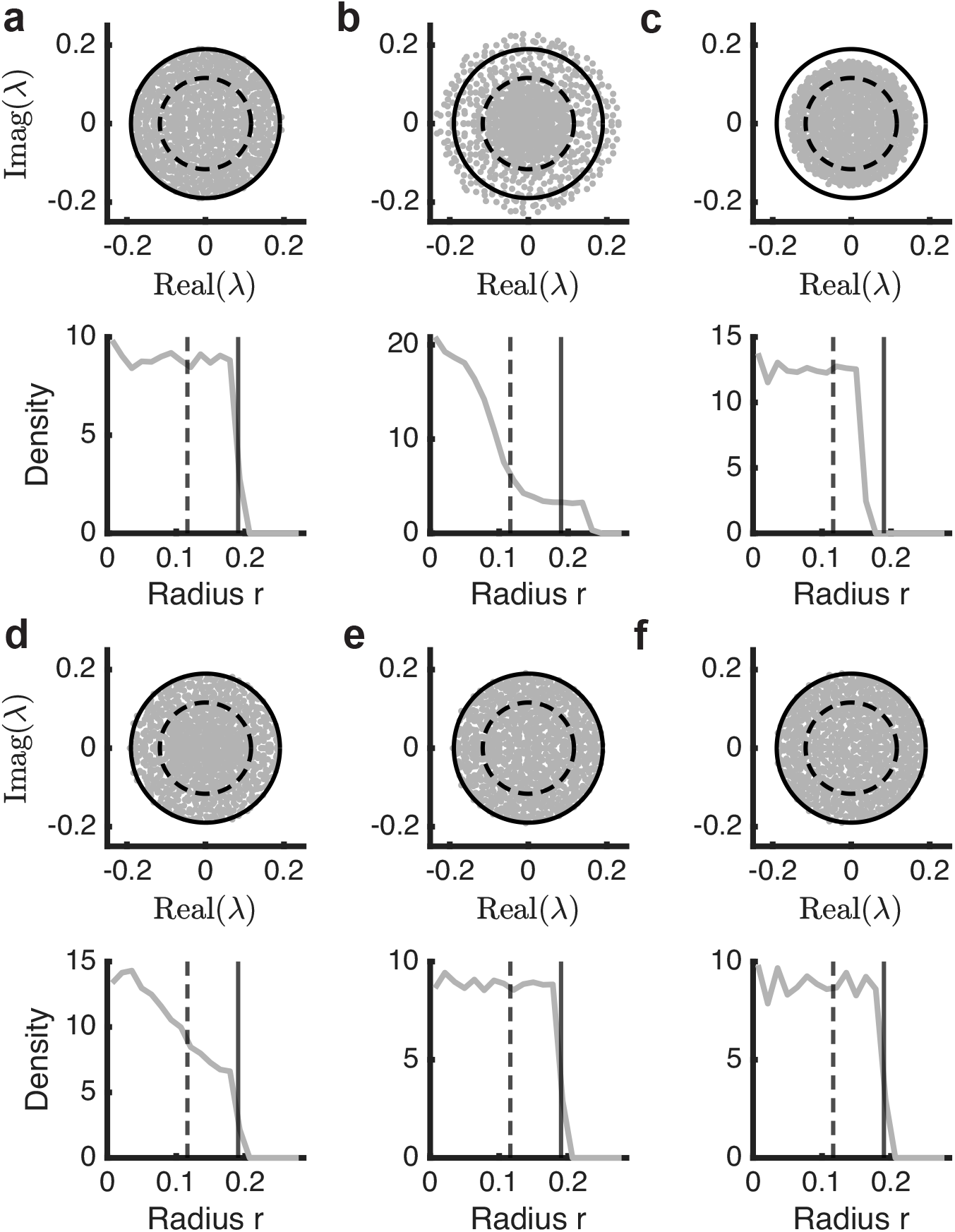
Sampled eigenvalues of various connectivity matrices on ℂ and the corresponding spectral density as a function of radial distance from (0,0). Solid black line – circle of radius *g*_0_ or support of the circular distribution. Dashed black line – circle of radius *g* with *κ*^2^ = 0.09. **a)** *W* with uniform variance (circular law). **b)** *W* with within-area structure. **c)** *W* with between-area structure. **d)** *W* with both within- and between-area structure. **e)** *W* with within-area structure in both areas. **f)** *W* with between-area structure across both areas.

**Figure S2.**
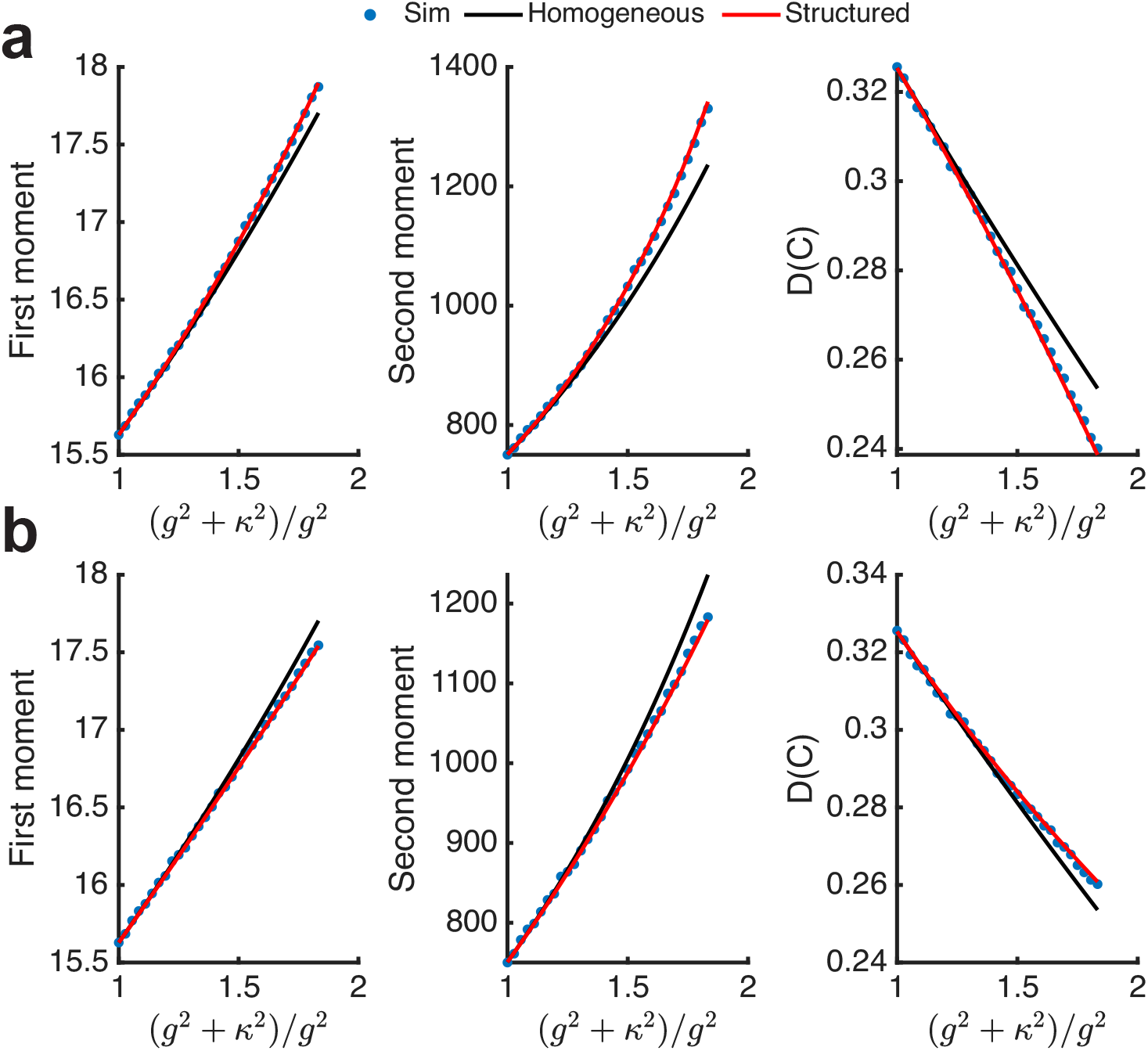
**a)** First moments, second moments, and *D*(*C*) across theory and sims after added within-area structure on area 2 without conserved variance. **b)** First moments, second moments, and *D*(*C*) across theory and sims after added between-area structure without conserved variance.

**Figure S3.**
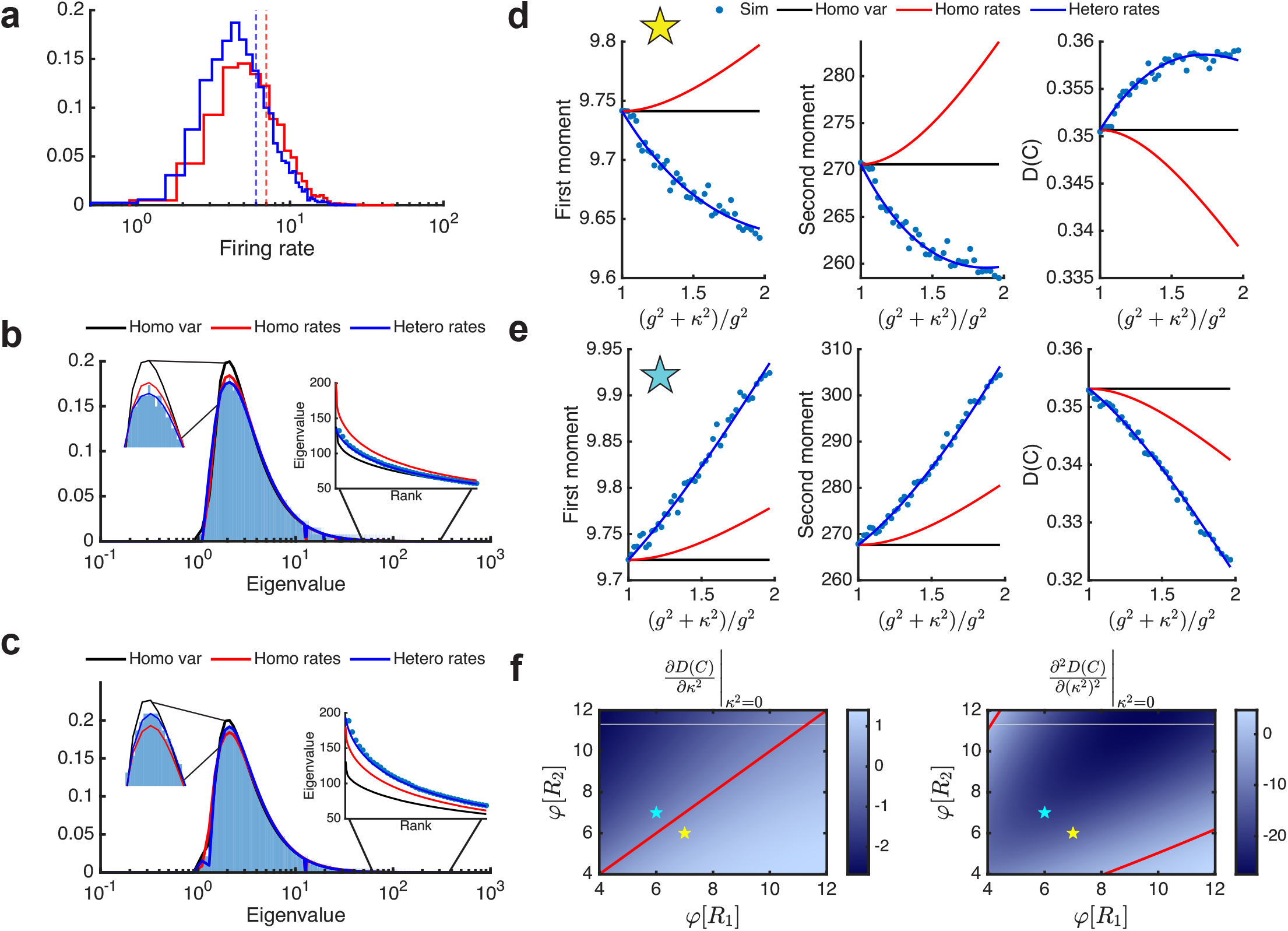
**a)** Two different log-normal firing rate distributions *R*_1_ and *R*_2_. In this example, *φ*[*R*_*i*_] = 6, 7 for *i*∈{1, 2}with area-specific *D*(*R*) fixed at 0.8. **b)** Simulation and theory agreement of the spectral density and its tail after within-area structures on area 2, when area 2 has a lower average firing rate. **c)** Simulation and theory agreement of the eigenvalue distribution and its tail after within-area structures on area 2, when area 2 has a higher average firing rate. **d)** Theory and simulations for the moments and *D*(*C*) match for added connection variance on area 2 when area 2 has the lower average firing rate. **e)** Theory and simulations for the moments and *D*(*C*) match for added connection variance on area 2 when area 2 has the higher average firing rate. **f)** Heatmaps of the coefficients of the Taylor’s expansion of *D*(*C*) about *κ*^2^ = 0. Left: 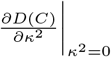 positive when area 2 has a lower mean firing rate, and negative when area 2 has a higher mean firing rate. Right: 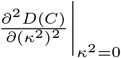 is always negative in the two examples shown. Red lines depict when values are zero.

**Figure S4.**
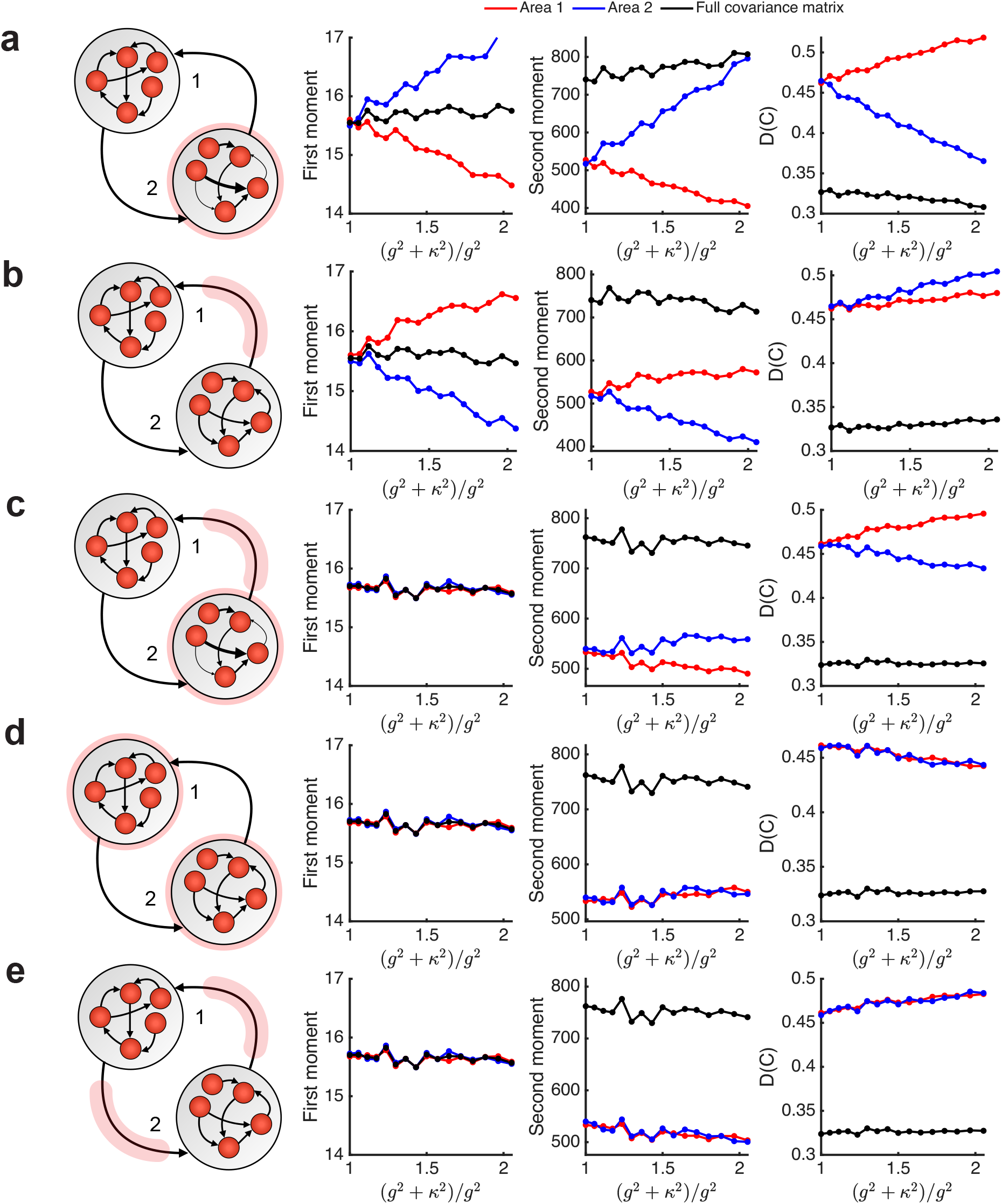
First moments, second moments, and *D*(*C*) for each area and the full covariance matrix across various structures. **a)** Within-area structure in area 2. **b)** Between-area structure from area 2 to 1. **c)** Both within- and between-area structure on area 2. **d)** within-area structures in both areas. **e)** between-area structures in both areas.

